# Structural and functional analysis of the *Mycobacterium tuberculosis* MmpS5L5 efflux pump presages a pathway to increased bedaquiline resistance

**DOI:** 10.1101/2025.06.24.661325

**Authors:** Adam J. Fountain, Jan Böhning, Stephen H. McLaughlin, Tomos E. Morgan, Paul H. Edelstein, Mark Troll, Meindert H. Lamers, Tanmay A. M. Bharat, Ben F. Luisi, Lalita Ramakrishnan

## Abstract

Bedaquiline, an antitubercular drug that targets ATP-synthase, is a key component of a new oral drug regimen that has revolutionized the treatment of multi drug resistant tuberculosis. Clinical bedaquiline resistance in *Mycobacterium tuberculosis* has rapidly emerged, primarily due to mutations in the transcriptional repressor, *Rv0678* that result in upregulation of the Resistance-Nodulation-Division (RND) efflux pump MmpS5/MmpL5 (MmpS5L5). Here, to understand how MmpS5L5 effluxes bedaquiline, we determined the structure of the MmpS5L5 complex using cryo-electron microscopy, revealing a novel trimeric architecture distinct from the canonical tripartite RND efflux pumps of Gram-negative bacteria. Structure prediction modelling in conjunction with functional genetic analysis indicates that it uses a periplasmic coiled-coil tube to transport molecules across the cell wall. Structure-guided genetic approaches identify MmpL5 mutations that alter bedaquiline transport; these mutations converge on a region in MmpL5 located in the lower portion of the periplasmic cavity, proximal to the outer leaflet of the inner membrane, suggesting a route for bedaquiline entry into the pump. While currently known clinical resistance to bedaquiline is due to pump upregulation, our findings that several MmpL5 variants increase bedaquiline efflux may presage the emergence of additional modes of clinical resistance.

**Significance Statement:** Resistance to bedaquiline, a cornerstone drug for treating multidrug-resistant tuberculosis, is rapidly emerging due to mutations that upregulate expression of the MmpS5L5 efflux pump. Here, we reveal the cryo-EM structure of this pump, showing a novel trimeric architecture and a unique α-helical coiled-coil tube for drug transport. Structure-guided genetic analysis identifies MmpL5 variants that further increase bedaquiline efflux, suggesting potential resistance mechanisms beyond pump upregulation.

## Introduction

Multidrug resistance is a major obstacle for the effective treatment of tuberculosis, with an estimated 400,000 cases of MDR-TB each year (1). Bedaquiline underpins the efficacy of the BPaL(M) regimens which have significantly improved the ability to treat MDR-TB (2,3). Since its implementation, however, bedaquiline resistance has emerged at an alarming rate (4–6). Bedaquiline resistance is associated with increased rates of treatment failure (7–11) and threatens the future efficacy of these regimens. Clinical bedaquiline resistance is primarily caused by mutations in the transcriptional repressor *Rv0678* which result in upregulation of the adjacent *MmpS5L5* operon (12–14). MmpS5 and MmpL5 form a complex that functions as a multidrug efflux pump (*SI Appendix* Fig. S1) which confers resistance to bedaquiline (12,13) and other hydrophobic drugs (14–17) as well as exporting the iron-scavenging siderophores mycobactin and carboxymycobactin (18,19). MmpS5 is a small (∼15 kDa) single helix transmembrane (TM) protein containing a C-terminal immunoglobulin-like (Ig-like) domain (18,20). MmpS5’s operonic partner, MmpL5, is a 12-TM Hydrophobe-Amphiphile-Efflux 2 (HAE-2) subfamily RND transporter (21) that is evolutionarily divergent from the well-characterised Gram-negative HAE-1 efflux pumps (22). Structures of other HAE-2 family MmpL transporters are monomeric (23–27); however, TIRF photobleaching experiments have shown that MmpL5 exists as a trimer in vivo, and that this is likely the functional assembly in the bacterium (28). Despite its critical role in conferring bedaquiline resistance, the architecture of the MmpS5L5 complex and whether it forms a tripartite assembly similar to Gram-negative efflux pumps (29) remains unclear. Moreover, the specific mechanisms by which MmpS5L5 recognizes and transports bedaquiline are unknown. Here, we integrate single-particle cryo-EM, protein structure prediction, and genetic approaches to determine the architecture of the MmpS5L5 assembly and understand its mechanism of drug efflux and substrate recognition.

## Results

### MmpS5L5 forms a trimeric efflux pump assembly

To determine the architecture of the MmpS5L5 complex, we purified *M. tuberculosis* MmpS5 and MmpL5 Δ494-688 overexpressed in *M. smegmatis* (*Msm*) by affinity purification using a C-terminal GFP-FLAG tag on MmpL5 followed by size exclusion chromatography (*SI Appendix* Fig. S2). SDS-PAGE and anti-MmpS5 immunoblots of peak fractions showed that MmpS5 co-purified with MmpL5 (*SI Appendix.* Fig. S2*C*). Cryo-EM single particle analysis (Materials and methods) revealed both monomeric MmpL5 and a subset of trimeric MmpS5L5 particles. To address preferred orientation, data were collected at 0°, 20°, and 40° tilts, yielding 3.2 Å maps of both monomeric MmpL5 and trimeric MmpS5L5 (Fig. 1*A*). Our structure reveals that the MmpS5L5 complex is a C3 symmetric homotrimer. MmpS5’s single transmembrane helix interacts with MmpL5 helices TM5 and TM8; however, density for the C-terminal Ig-like domain is absent in the map.

**Figure 1:**
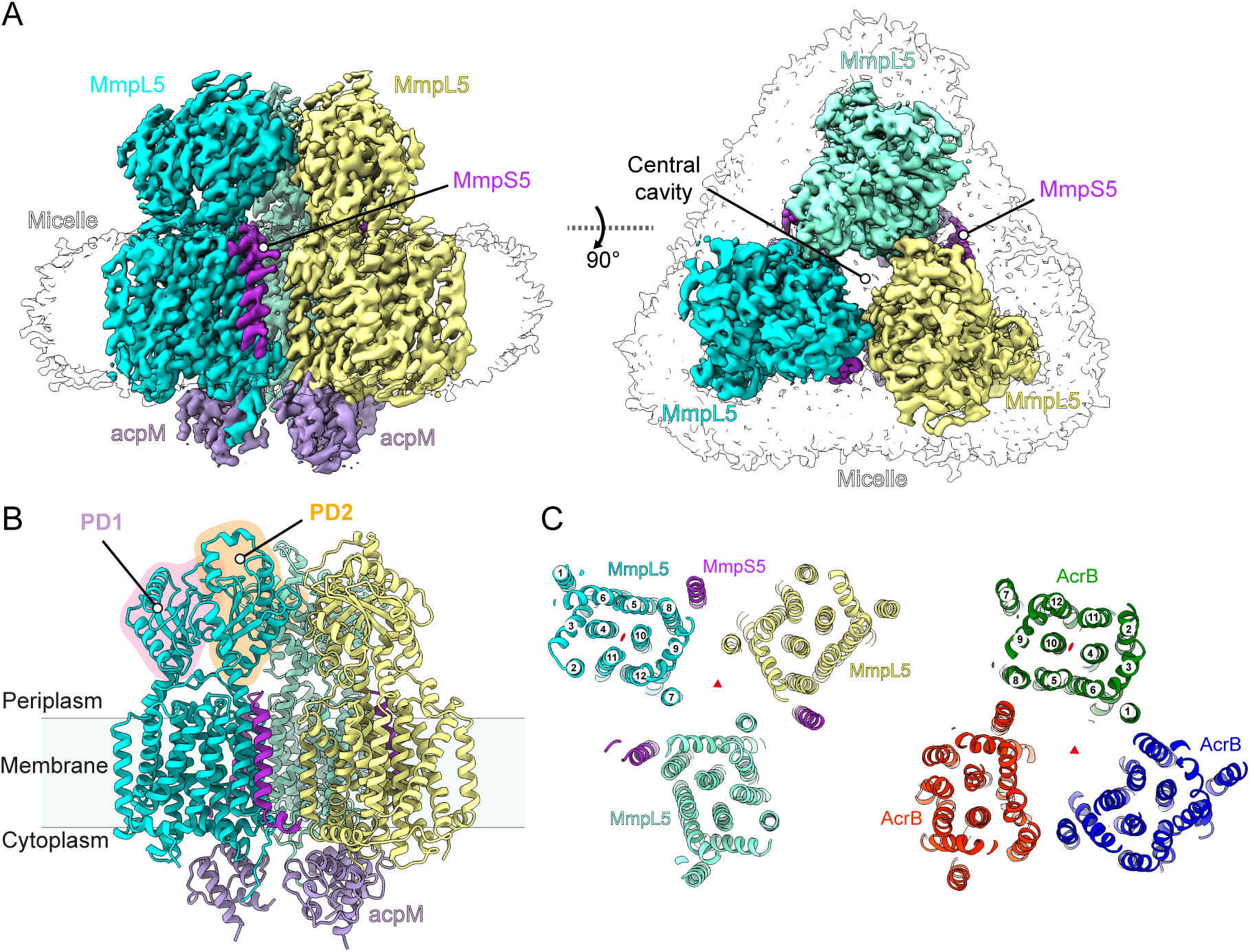
Architecture of the trimeric MmpS5L5-AcpM complex. (A) Cryo-EM density map of the MmpS5-MmpL5ΔCC-AcpM complex. (B) Structural model of MmpS5-MmpL5ΔCC-AcpM complex. Periplasmic domain PD1 (Pink) and PD2 (Orange) highlighted (C) Slice view of the transmembrane domains of MmpS5L5 (left) and AcrB (right). TM helix numbers are indicated.

MmpL5 shares its 12-TM helix RND fold with Gram-negative transporters such as AcrB. This fold exhibits two-fold pseudo-symmetry between the two 6-TM helical bundles (Fig. 1*C*). MmpL5 possesses two periplasmic α+β sandwich fold domains (PD1 and PD2, respectively), with one present on top of each 6-TM bundle. In contrast, AcrB contains two stacked porter domains with an additional docking domain per 6-TM bundle (*SI Appendix*. Fig. S6). The trimeric organisation of MmpS5L5 differs from its Gram-negative RND counterpart, with protomer interactions occurring across different faces of the TMDs (Fig. 1*C*). In AcrB, TMs 1 and 8 mediate the interaction between protomers within the transmembrane domain, with TM helices 3,5,6 and 8 lining the internal cavity. In MmpL5, TM7 and TM12 in adjacent protomers are bridged by lipid-like densities, with TM helices 7,8,9 and 12 lining the internal cavity. These data show that MmpS5L5 is a novel trimeric efflux pump assembly, whose architecture is distinct from the tripartite efflux assemblies from Gram-negative bacteria.

Remarkably, few direct protein-protein contacts are observed between MmpL5 protomers within the trimer. The only major interaction identified is in PD2, where Lys747 interacts with Phe473 and the carbonyls of Val475 in the adjacent MmpL5 protomer’s PD2. Rather than direct protein-protein contacts, we find lipid-like densities at the interfaces between subunits within the TMD (*SI Appendix* Fig. S7*A*). Density that we model as a phospholipid interacts with MmpS5 Trp9 via a classic lipid-protein interaction motif (30), where the indole nitrogen of tryptophan forms a hydrogen bond with the ester group of the nearby phospholipid. The acyl chain of the phospholipid interacts via hydrophobic interactions with the TMs of both MmpS5 and MmpL5. Weak densities resembling ordered acyl chains are seen in the internal cavity (*SI Appendix*. Fig. S7*A*), hinting that this region is also occupied by lipids, similar to what is observed in AcrB (31). These lipid-mediated contacts may explain the complex’s lability in detergent and strongly suggest that lipids are essential for maintaining the trimeric assembly and function.

We observed density on the cytoplasmic face of MmpL5, which we assign as the endogenous *M. smegmatis* acyl carrier protein AcpM (MSMEG_4326) (Fig. 1*A*), as a similar density was observed at the corresponding location in the monomeric cryo-EM structure of the MmpL protein MSMEG_1382 from *M. smegmatis* (24). AcpM interacts with MmpL5’s basic cytoplasmic Iα2 helix and MmpL5’s unstructured C-terminus (residues 946–952), which forms an extended interface with AcpM (SI *Appendix*. Fig. 7*B*, right). Extending from AcpM, we observe density corresponding to Ser41 post-translationally modified with 4’-phosphopantetheine, which is sandwiched between two MmpL5 protomers, interacting with both Trp389 and Trp939. By bridging protomers using its phosphopantetheine moiety, AcpM binding likely contributes to the stability of the trimeric assembly of MmpS5L5.

### MmpL5’s coiled-coil domain forms a periplasmic tube

Our cryo-EM structure resolved the core transmembrane and periplasmic domains but lacked MmpL5’s unique 196 amino acid insertion within periplasmic domain 2 (PD2), which is predicted to form an extended anti-parallel coiled-coil with a disulfide bond at the tip. To understand the contribution of this region to MmpL5 function, we first confirmed that purified MmpL5 (Residues 493–682) has coiled-coil secondary structure by circular dichroism spectroscopy (*SI Appendix* Fig. S9*D,E*). Intact mass spectrometry of the purified coiled-coil domain shows a +2 Da shift after reduction by TCEP, indicating that an intramolecular disulfide bond forms between C591 and C597 (*SI Appendix* Fig. S9*F*). Genetic deletion of the coiled-coil domain abolished the ability to complement the Δ*S5L5* strain for all tested drugs (*SI Appendix* Fig. S9*B,C*), showing that the coiled-coil domain is necessary for efflux. As substrates such as siderophores must be exported to the extracellular environment, we speculated this domain might associate to form a periplasmic conduit for transport. To gain insight into the full-length complex including the coiled-coil domain, we used AlphaFold2 to model a trimeric MmpS5L5 complex. This produced a high-confidence prediction with an average pLDDT score of 84.4, pTM of 0.84 and ipTM of 0.84 (*SI Appendix* Fig. S8*A*) agreeing well with the experimental structure (RMSD = 1.1 Å) (*SI Appendix* Fig. S8*C*).

In the AlphaFold2 model, the coiled-coils interact to form an extended (∼ 13 nm) hexameric alpha-helical barrel with a 7–9 Å diameter lumen lined with methionine residues (Fig. 2*D*). The coiled-coil domain contains multiple type I and type II heptad (*gabcdef*) repeats (32,33) (Fig. 2*C*, *SI Appendix* Fig. S9). Analysis of the geometry of the predicted tube showed “extended” knobs-in-holes packing (Fig. 2*C*) that is consistent with the sequence-structure relationship defined for de novo designed alpha-helical barrels (33–35). As small molecules can be accommodated within alpha-helical barrels (36), we suggest that MmpL5’s substrates transit through the lumen of this tube. The side chain of methionine is highly flexible, mildly hydrophobic, and the thioether group can make S/π interactions with aromatic moieties (37,38), and we speculate that these characteristics allow varied aromatic/lipophilic substances to transit along the lumen.

**Figure 2:**
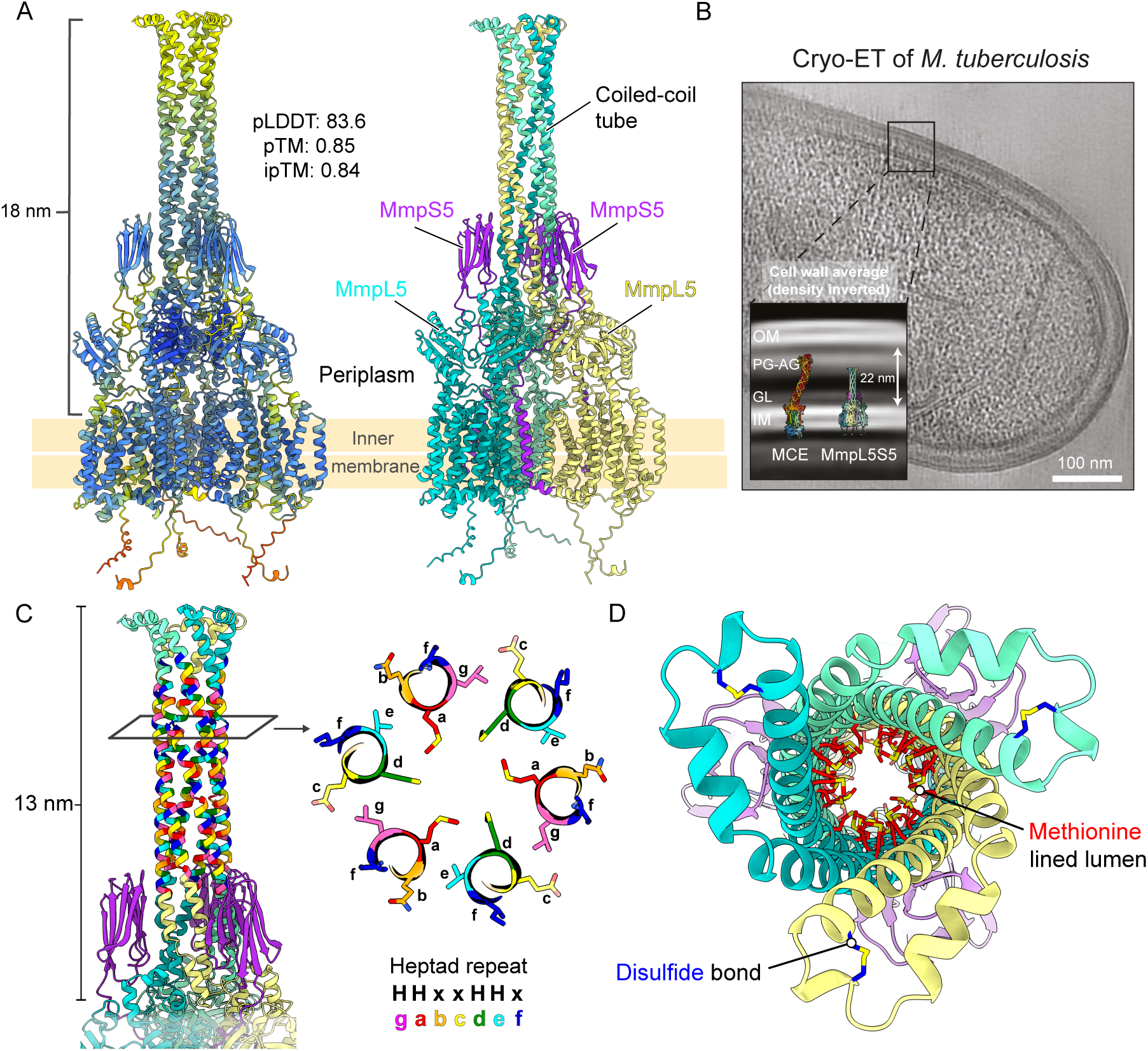
MmpL5’s coiled-coil domain forms a 13 nm alpha-helical tube. (A) Structural model of the AlphaFold2 predicted structure of MmpS5L5 trimers, coloured according to per-residue pLDDT value (left) and by subunit (right). (B) Tomogram of *M.tuberculosis* mc^2^6206. Inset, averaged cell wall density with the AlphaFold2 model of MmpS5L5 and the experimentally determined MCE1 complex (PDB: 8FEF (93)) overlaid with their transmembrane domains on the inner membrane. IM – Inner membrane, GL – Granular layer, PG/AG – Peptidoglycan/Arabinogalactan, MM – Mycomembrane. (C) AlphaFold2 model of MmpL5’s coiled-coil domain, with residues coloured according to position in the heptad repeat (*gabcdef*). (D) Top view down the axis of the tube. The interior methionine residues are highlighted in red. A conserved disulfide bond is coloured blue.

The distance from MmpL5’s transmembrane domain to the tip of the coiled-coil is ∼18 nm. Averaged cell wall density from electron cryotomography (cryo-ET) of *M. tuberculosis* shows that the length of the coiled-coil is insufficient to cross the periplasm and mycomembrane and extends as far as the peptidoglycan-arabinogalactan layer (Fig. 2*B*). This indicates that substrates must traverse the remaining periplasmic space and the mycomembrane via a yet unidentified mechanism to reach the extracellular environment.

### MmpS5 is required for efflux activity

We next focused on the role of MmpS5 in the overall assembly and function. In our AlphaFold2 prediction of trimeric MmpS5L5, MmpS5 bridges across all three MmpL5 subunits: MmpS5’s TM domain interacts with MmpL5’s TM8, whilst MmpS5’s Ig-like domain spans the neighbouring two MmpL5 protomers, wrapping around to interact across the seam formed between coiled-coil domains (Fig. 3*A*). This organisation explains the observation that MmpS5 is required for MmpL5 trimerization (28). The interaction is likely to be specific and functionally important for transport activity, as MmpL5 alone is unable to complement the ΔS5L5 mutant (Fig. 3*B*) (28), and other MmpS paralogs are unable to substitute for MmpS5 (Fig. 3*B*). Amino acid substitutions that disrupt the predicted MmpS5-MmpL5 interaction interface reduce or abolish pump activity: We find that MmpS5 C137S mutations that prevent the formation of a disulfide bond (20) in MmpS5’s Ig-like domain eliminate function (*SI Appendix* Fig. S10). Likewise, we find that K54A and K140A substitutions that disrupt salt bridges between MmpS5 and MmpL5 reduce function. Together, these data point to a specific requirement for MmpS5 and that a direct interaction between MmpS5 and MmpL5 is required for pump function.

**Figure 3:**
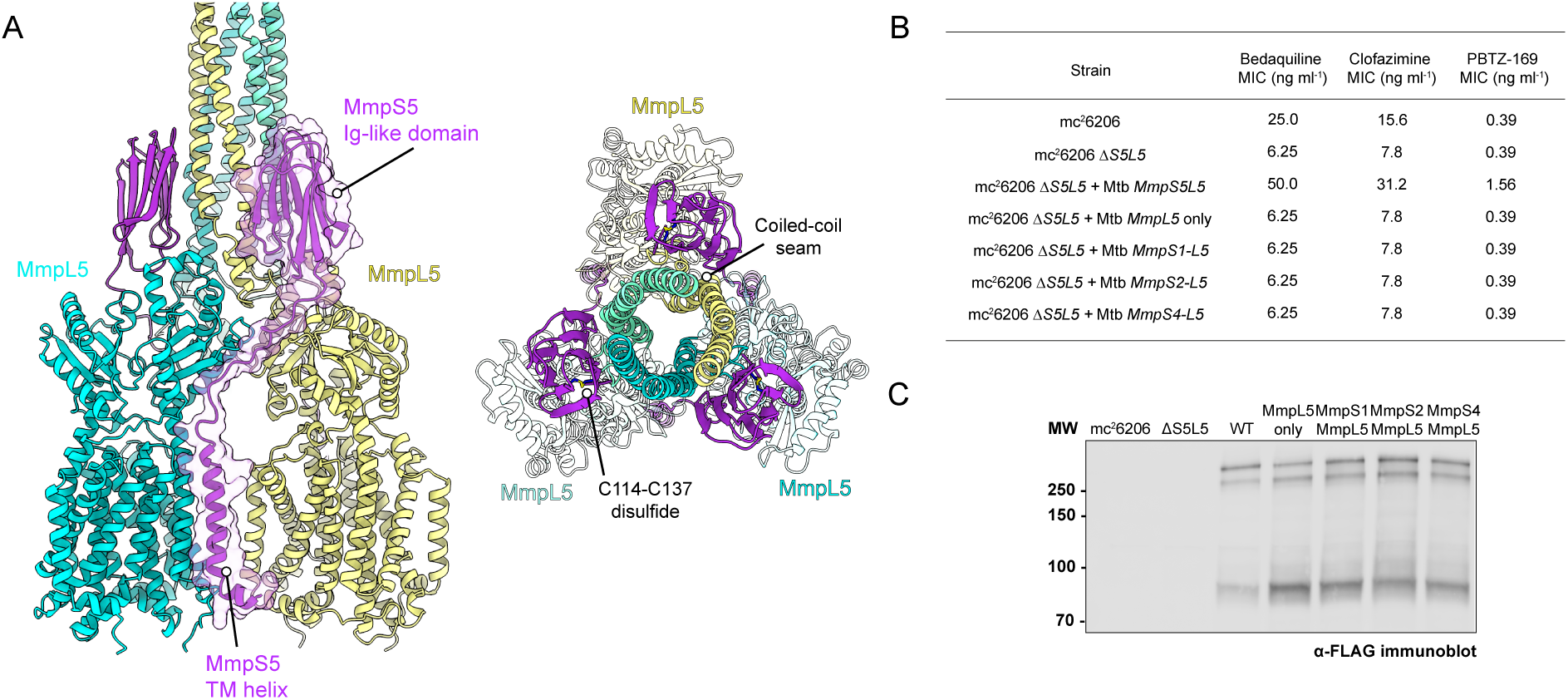
MmpS5 spans three MmpL5 protomers and is essential for efflux activity. (A) MmpS5’s TM interacts with TM helix 8 of MmpL5. Beta strand 2,3,5 and 8 of MmpS5’s Ig-like domain interacts across the interface formed between (B) MIC values of Bedaquiline, Clofazimine and PBTZ-169 show that MmpS5 is necessary for MmpL5 to efflux drugs. Paralogs MmpS1, MmpS2 and MmpS4 are unable to compensate for MmpS5. (C) Anti-FLAG immunoblot showing MmpL5 expression in all strains.

### Substrate recognition and bedaquiline efflux

In vitro, bedaquiline binds purified *Mtb* MmpL5 with a K_d_ of 2.2 µM (Fig. 4*A*, *SI Appendix* Fig. S*11*), as measured by surface plasmon resonance (SPR). Despite incubation with 50 µM (∼25 × K_d_) bedaquiline prior to cryo-EM specimen preparation, no clear density corresponding to bedaquiline was observed. As we were unable to identify a binding site for bedaquiline by cryo-EM, we turned to genetics and comparative sequence analysis to identify regions of MmpL5 that are important for substrate recognition or transport. We exploited the observation that the MmpS5L5 paralog, MmpS4L4, which shares the ability to export the iron siderophore mycobactin (18), fails to rescue the bedaquiline sensitivity of the Δ*S5L5* strain despite high sequence and structural similarity (Fig. 4*B*). RND transporters typically recognise their substrates in a cavity formed between their extracellular domains (39,40). Therefore, we examined amino acid differences in the periplasmic domains of MmpL4 and MmpL5 and selected those substitutions not found in a multiple sequence alignment of MmpL5 proteins, indicating possible functional importance (*SI Appendix* Fig. S12). When introduced into MmpL5, N444K and Q196M reduce the bedaquiline MIC 4-fold relative to wild-type MmpL5. Q196M, however, has no effect on the clofazimine MIC and increases the PBTZ-169 MIC 2-fold. The effects of the mutations are not due to reduced protein expression, confirmed by immunoblot (*SI Appendix* Fig.16*B*), or protein inactivation, as this would be expected to reduce the MICs of all drugs. Expanding this paralog-guided mutagenesis strategy to MmpS1L1 and MmpS2L2 we identified Q150V, Q196R, D417L, H832Y and N444E as activity variants in MmpL5 (Fig. 4*C* and *D, SI Appendix* Fig. S12)

**Figure 4:**
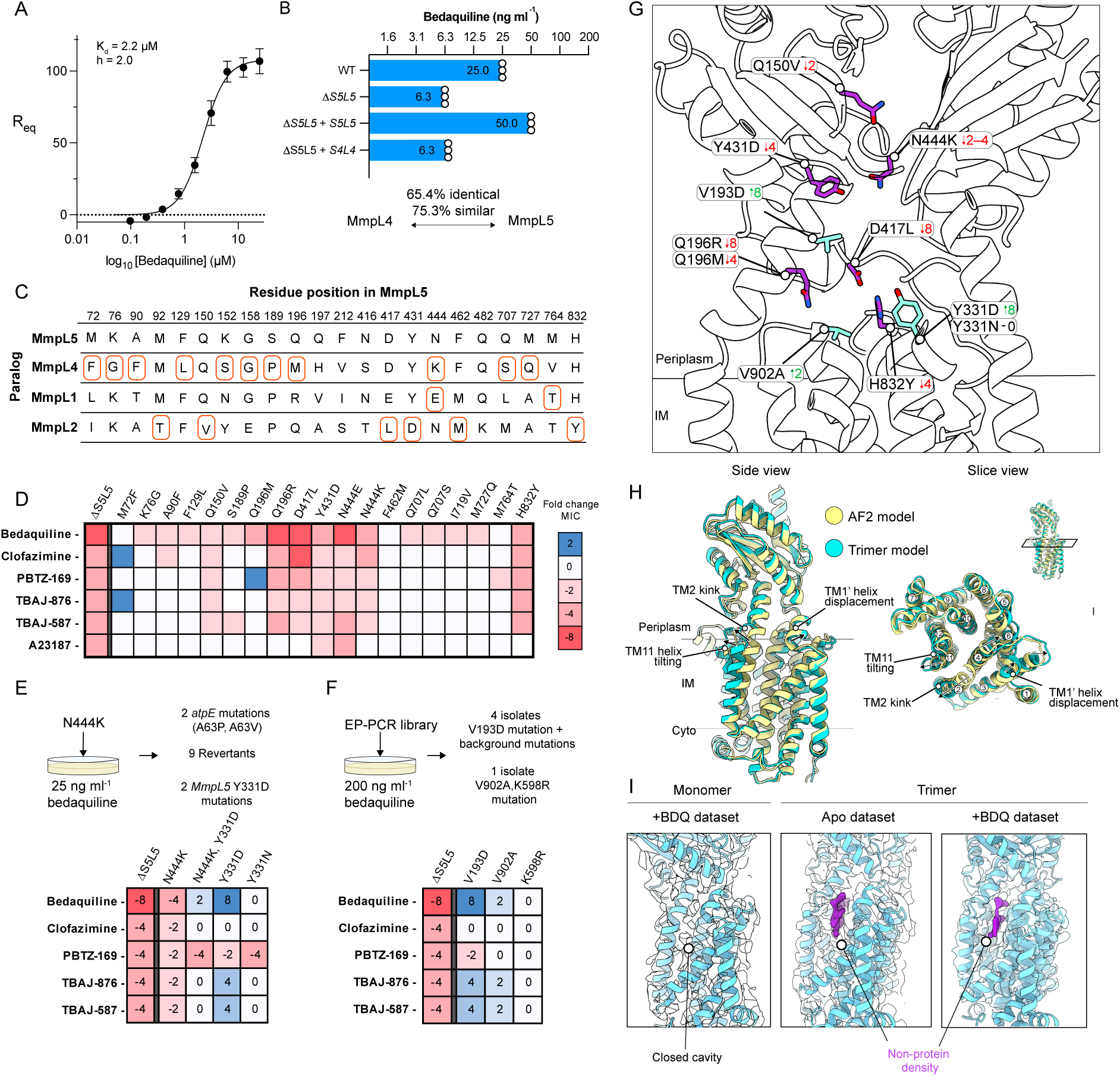
MmpL5 activity variants converge on a region of MmpL5 in a lower portion of the periplasmic cavity. (A) Plot of R_eq_ against substrate concentration on a log_10_ axis. Points represent the mean value of three technical replicates. Error bars represent standard deviation. K_d_ values and Hill coefficient (h) were calculated using a specific binding with Hill slope model. (B) Bar chart showing the Bedaquiline MIC values for the indicated strains. Open circles indicate technical replicates. (C) Alignment of MmpL5/4/2/1 at the selected positions. Amino acids for each paralog are shown. Amino acid changes chosen for mutation are indicated in red boxes. (D) MIC values for the selected amino acid substitutions are coloured according to fold change in MIC versus the complemented MmpS5L5 strain. (E–F) Upper, Summary diagram showing strategy for isolating suppressor/activity variants. (G) Experimental structural model of MmpL5 indicating residues that alter bedaquiline transport. Fold changes are given alongside the indicated substitution. Green indicates positive fold change in bedaquiline MIC, red indicates a negative fold change in bedaquiline MIC. (H) Overlay of a protomer from MmpS5L5 trimer structure with the AlphaFold2 model of MmpL5. Arrows indicate displacement of MmpL5 relative to the AlphaFold model. (I) A non-protein density in the TM1–4 cleft of MmpL5 that is present in both ‘apo’ and bedaquiline incubated samples. No density is observed in this cleft in the monomer structure.

**Figure 5:**
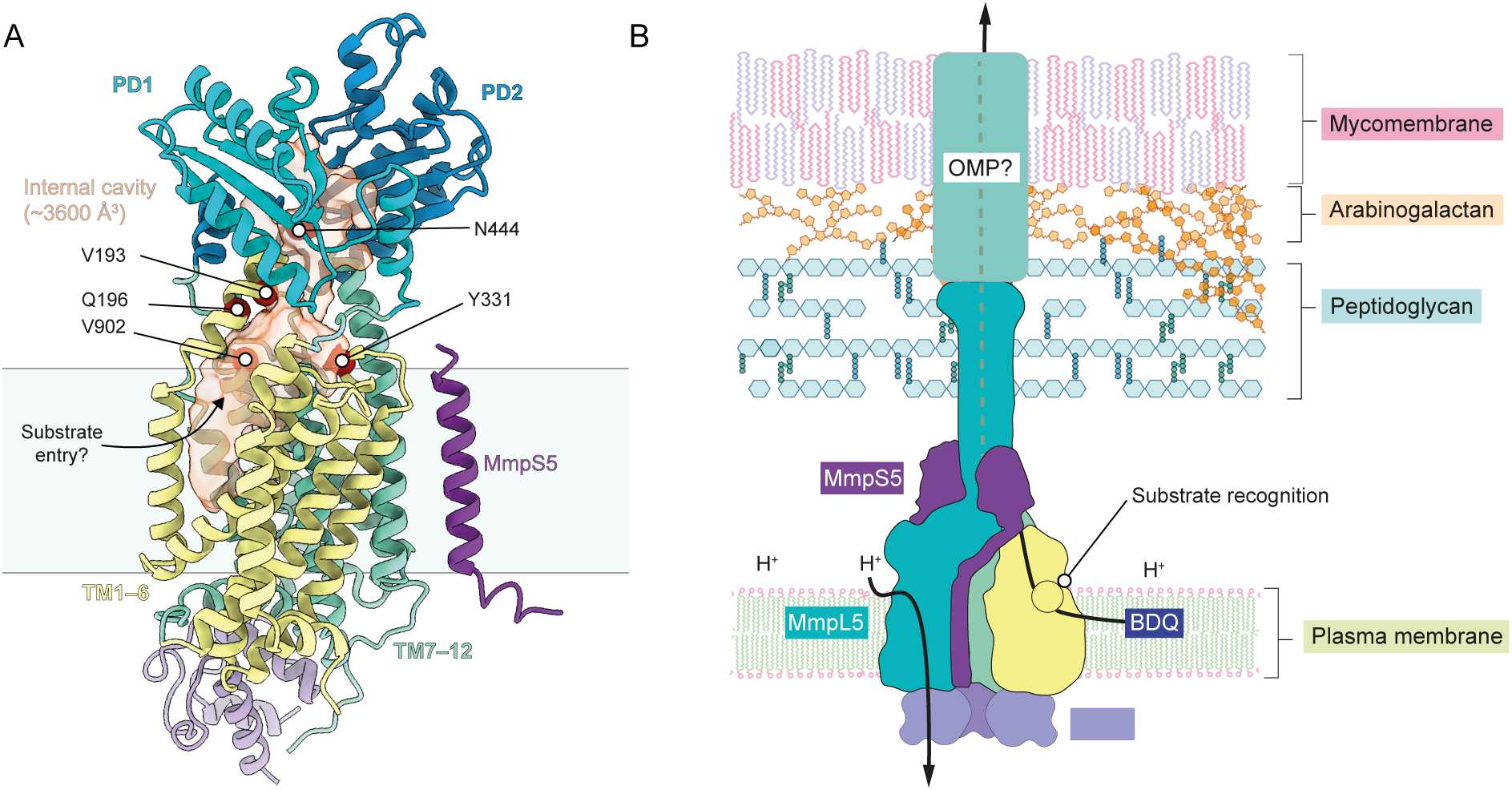
Model for bedaquiline entry and transport. (A) Experimental structural model of MmpL5, with residues that alter the activity of MmpS5L5 indicated. PD1/PD2 – Periplasmic domain 1 or 2 respectively. Residues identified as being functionally important for bedaquiline efflux are indicated with red spheres. (B) The model for MmpS5L5 mediated efflux of bedaquiline and other hydrophobic anti-tubercular drugs. Hydrophobic drugs like bedaquiline partition from the membrane into the periplasmic cavity of MmpL5 via TMs 1–4. The essentiality of the coiled-coil domain and its unique methionine lined lumen, suggest that substrates are transported via this conduit. The length of this domain is too short to cross the entire cell wall. Strong sequence conservation of a hydrophobic surface at the tip of the coiled-coil domain suggest that an unknown outer membrane factor facilitates further substrate transport across the mycomembrane.

To further probe these functionally important residues, we used a *M. marinum mmpL5*::Tn (MMAR_1005::Tn) strain complemented with *Mtb MmpS5L5* as a surrogate to speed up genetic analysis. *M. marinum* has a conserved *Rv0678*-*MmpS5L5* locus and exhibits the same pattern of bedaquiline sensitivity as *M. tuberculosis* with knockout and complementation (*SI Appendix* Fig. S13). We then isolated intragenic suppressor mutations of the sensitising N444K mutation by plating ∼ 3 × 10^9^ CFU of Mm mmpL5::Tn + MmpS5L5 N444K on 25 ng ml^-1^ bedaquiline. Screening resistant *M. marinum* isolates identified a secondary Y331D substitution in MmpL5 that restored MIC to wild-type levels. We incorporated the Y331D mutation into the wild-type and N444K MmpL5 protein sequence and re-introduced the resulting construct into a clean *Mtb* Δ*S5L5* background. N444K, Y331D double mutants restored the bedaquiline MIC to 2-fold higher than wild-type levels (Fig. 4*E*), showing Y331D is an intragenic suppressor of the N444K allele. Unexpectedly, Y331D alone increased the bedaquiline MIC 8-fold above the wild-type MmpS5L5. Moreover, the increased resistance appears to be specific for bedaquiline, since the MIC of clofazimine is unaltered by the Y331D mutation (Fig. 4*E*). We ruled out increased protein expression as the cause of the increased bedaquiline resistance of the Y331D allele (*SI Appendix* Fig.16*C*). Y331 is located at the top of TM helix 6, at the bottom of a cavity formed by the two periplasmic domains and is adjacent to Q196 (Fig. 4*G*). The enhancement of bedaquiline efflux by Y331D mutation may be by electrostatic interaction with the positively charged dimethylamino group of bedaquiline, as Y331N mutations did not exhibit the same increase in MIC (Fig. 4*E*). This suggests that charge-charge interactions are important for bedaquiline recognition, as has been seen for the interaction of cationic substrates in other efflux pumps (41). Long-range electrostatic interactions can facilitate molecular recognition above the diffusion limit (42), potentially providing an explanation for the increased activity.

To identify other mutants that increase bedaquiline transport activity, we created a library containing MmpS5L5 mutants produced using low-fidelity (‘error-prone’) PCR (*SI Appendix* Fig. S14). This library was transformed into *M. marinum* mmpL5::Tn and plated on 2× MIC bedaquiline (0.2 µg ml^-1^) to select for increased pump activity. Four out of five bedaquiline-resistant isolates contained a V193D mutation along with differing background mutations (Fig. 4*F*, *SI Appendix* Fig. S15). Reintroduction of *MmpS5L5* V193D into a clean *Mtb* Δ*S5L5* background increased the bedaquiline MIC 8-fold over wild-type MmpS5L5; however, V193D unexpectedly reduced the MIC for PBTZ-169 and did not impact clofazimine MIC (Fig. 4*F*). V193D mutations also increase MICs of the bedaquiline derivatives TBAJ-587 and TBAJ-876 4-fold (Fig. 4*F*). The V193D mutation did not alter MmpL5 expression (*SI Appendix* Fig. 16*D*). We also identified an isolate with a V902A, K598R double mutation; when tested individually the V902A variant increases the bedaquiline MIC 2-fold (Fig. 4*F*). V193 is located in TM helix 2 and V902A at the top of TM helix 12, adjacent to both Q196 and Y331 (Fig. 4*G*). The convergence of two independent methods on the same region in the lower portion of the periplasmic cavity of MmpL5 strongly supports this region being functionally important for the efflux of bedaquiline (Fig. 4*G*) and we propose that this is the initial site of recognition.

Further supporting our proposal that this is a substrate binding site, in our experimental trimer structure, TM2 is kinked and TM11–12 are displaced outwards compared to both the monomeric and AlphaFold2 models, significantly increasing the volume of the hydrophobic TM1–4 cavity (Fig. 4*H*). A distinct non-protein density is clearly visible within this cavity in both ‘apo’ and bedaquiline-saturated trimeric structures (Fig. 4*I*), suggesting that molecules can enter this cavity. Crucially, the monomeric MmpL5 structure derived from the same cryo-EM data lacks this kinked TM2 helix and the putative substrate density (Fig. 4*I*). This suggests that trimerization induces a specific conformational change that opens this entry cavity, potentially facilitating substrate access from the inner membrane leaflet. As bedaquiline is highly lipophilic (logP ∼ 7.2), it is plausible that the drug can partition out of the plasma membrane via TMs 1–4 to reach this site within MmpL5. Substrate entry from the membrane via TMs 1–4 is observed for other RND family transporters (43,44) and is similar to the membrane-access ‘CH1’ channel seen in AcrB (45,46).

## Discussion

Bedaquiline resistance caused by upregulation of MmpS5L5 in *M. tuberculosis* is a rapidly growing threat to the efficacy of MDR-TB treatment regimens (47). Our structure shows that MmpS5L5 is a novel class of trimeric RND efflux pump and genetic analysis allows us to propose a model for substrate transport. MmpS-MmpL loci are found in many species of clinically important tuberculous and non-tuberculous mycobacteria and are associated with resistance to a wide variety of antimicrobial agents (48,49) and so our work provides a model for understanding this important protein family. The structural difference between the trimeric Gram-negative and mycobacterial RND assemblies suggests independent evolution of a trimeric organisation to facilitate transport across the mycobacterial cell wall.

Our combined structural and genetic approaches identified a site within the lower portion of MmpL5’s periplasmic cavity as functionally critical for bedaquiline efflux, which we propose is the initial binding site for substrates. Unlike the reported mycobactin binding site in MmpL4 (23), our data suggest bedaquiline recognition occurs in a distinct protein region. How substrates move from this region to enter the coiled-coil domain is unclear and would likely require a series of conformational changes. AcrB uses coupled conformational cycling between protomers (50,51) known as “functional rotation” to efflux substrates. Given its trimeric organization, MmpS5L5 may employ ‘functional rotation’ similar to AcrB, a mechanism that may be advantageous as it can enhance efflux rate (52).

The observation that MmpL5’s coiled-coil domain is too short to span the periplasm and mycomembrane leads to the question of how substrates are exported through the hydrophobic mycomembrane. In Gram-negative bacteria, transport across the outer membrane is facilitated by the outer membrane channel TolC (53–55). Mycobacteria, which are considered ‘neo-diderm’, share none of the OM machinery – including *TolC* – with Gram-negative bacteria, having independently evolved an outer membrane from the otherwise monoderm Actinobacteria (56). Strong sequence conservation of a hydrophobic surface at the tip of MmpL5’s coiled-coil domain (*SI Appendix* Fig. S9*G*-*H*), suggests an unknown protein may interface with MmpL5 to facilitate substrate transport across the mycomembrane, strengthening previous proposals for an as-yet unidentified mycomembrane partner (18,28,57).

No mutations in MmpS5 or MmpL5 are currently linked to clinical bedaquiline resistance (58). Our genetic studies identified Y331D, V902A, and V193D mutations that increase the activity of MmpL5, demonstrating that MmpL5 has “evolutionary space” to increase efflux of bedaquiline. These variants are absent in all publicly available genomes from clinical *Mtb* isolates (59); however, if *Rv0678* mutant strains were to circulate, and with continued bedaquiline selection pressure, these variants may be seen clinically. Indeed, AcrB substitutions in clinical isolates offer a precedent for drug transporter mutations that enhance resistance by altering substrate recognition (60,61). The rarity of bedaquiline-resistant *atpE* mutants in clinical isolates has led to the suggestion that *atpE* mutants are less fit *in vivo* (62), but not *in vitro* (63). MmpL5 mutations may provide an alternative path to high-level bedaquiline resistance without a strong fitness cost. While the bedaquiline derivatives TBAJ-876 and TBAJ-587 currently in Phase I and II clinical trials (64) have the potential to overcome *Rv0678*-mediated resistance because of their increased potency, our data show TBAJ-876 and TBAJ-587 are also vulnerable to resistance by MmpL5 activity mutations (Fig. 4*E* and *F*).

Bedaquiline is the cornerstone of treatment regimens for MDR-TB. Rapidly emerging bedaquiline resistance threatens the efficacy of this life-saving drug. We hope that elucidating the structure of the trimeric MmpS5L5 transporter and highlighting MmpL5 variants with increased bedaquiline resistance that our work will spur the urgent development of new bedaquiline derivatives that can avoid or overcome efflux by MmpS5L5, as well as the development of potent efflux pump inhibitors.

## Materials and methods

No statistical methods were used to predetermine sample size. The experiments were not randomized, and investigators were not blinded to allocation during experiments and outcome assessment.

### Bacterial strains

All *M. tuberculosis* strains used were derived from the double auxotrophic *M. tuberculosis* H37Rv mc^2^6206 Δ*leuCD* Δ*panCD* strain (65) (obtained from William Jacobs Jr.). Generation of *M. tuberculosis* H37Rv mc^2^6206 Δ*leuCD* Δ*panCD* Δ*mmpS5L5* strain was described previously (66). *M. tuberculosis* H37Rv mc^2^6206 and derivatives were cultured either on solid media on 7H10 + 10% (v/v) OADS agar plates supplemented with 50 μg ml^-1^ L-leucine and 24 μg ml^-1^ calcium pantothenate or in Middlebrook 7H9 broth supplemented with 10% v/v OADS, 0.2% glycerol, 0.05% Tween 80, 50 μg ml^-1^ L-leucine and 24 μg ml^-1^ calcium pantothenate, with 25 μg ml^-1^ kanamycin or 50 μg ml^-1^ hygromycin B as appropriate. The *M. marinum* mmpL5::Tn (MMAR_1005::Tn) strains used were derived from the *M. marinum* ‘M’ strain. *M. marinum* and *M.smegmatis* strains were grown in the same solid and liquid media as *M. tuberculosis*, but without L-leucine or pantothenic acid supplementation at 37 °C (*M. smegmatis*) or 33 °C (*M. marinum*).

### Compounds

The compounds used in MIC assays were as follows: Bedaquiline (Astatech #43211, Cas. No. 843663-66-1), Clofazimine (Toku-E #C201, Cas. No. 2030-63-9), PBTZ-169 (MedChemExpress #HY-12903, Cas. No. 1377239-83-2), OPC-167832 (MedChemExpress #HY-134940, Cas. No. 1883747-71-4), TBAJ-587 (MedChemExpress #HY-111747, Cas. No. 2252316-16-6), A23187 (Sigma-Aldrich #C7522, Cas. No. 52665-69-7), Nigericin (Sigma-Aldrich #SML1779, Cas. No. 28643-80-3). TBAJ-876 was a gift from Matt McNeil. All compounds were dissolved in anhydrous DMSO and stored as aliquots at −70°C except for nigericin which was dissolved in a 1:1 solution of DMSO and ethanol.

### DNA constructs

Plasmid constructs generated in this study are listed in table S1 and Genbank files for the plasmid DNA constructs are available at: https://doi.org/10.5281/zenodo.15720311. The oligonucleotides used to generate these constructs are listed in table S2.

### Complementation and expression plasmid construction

pFLAG-attP-Kan was constructed by in vivo assembly (IVA) cloning (67) by combining a fragment of pFLAG-attP (Addgene #110095) amplified using primer pair P298/P299 and the Kan cassette amplified from pMINTC3GH (Addgene #110077) using primer pair P300/P301. pMINTF3 was constructed by IVA cloning by combining a fragment of pFLAG-attP-Kan amplified using primer pair P721/P722 and the pMINT backbone from pMINTC3GH (Addgene #110077) using primer pair P723/P724. The pMEXC3GF expression vector was generated by replacement of the His tag of pMEXC3GH (Addgene #110082) with a FLAG tag using Q5 mutagenesis (NEB #E0554S) using primer pair P465/P466. Open reading frames were cloned into expression/complementation vectors by FX cloning as previously described (68,69).

### Construction of MmpS5L5 mutants

Mutations in MmpS5L5 containing single/double mutations were constructed by in-vivo assembly (67) using PCR-amplified pINIT-Cat *Mtb MmpS5L5* lacking the replacement region with ∼500 bp synthetic DNA fragments containing the desired sequence change (IDT eBlocks). For making MmpS5L5 deletions constructs, pINIT-Cat *Mtb MmpS5L5* was amplified with primers P366/P367 (Δ*MmpS5*), P368/P369 (ΔCoiled-coil), followed by DpnI digestion, phosphorylation with T4 polynucleotide kinase and blunt-end ligated with T4 DNA ligase. MmpS paralog-MmpL5 fusions were constructed using IVA cloning: pINIT-Cat *Mtb MmpS5L5* was amplified with primer pair P367/P583. MmpS paralogs with overhangs were amplified from Mtb H37Rv genomic DNA using primers P577/P578 (MmpS1), P579/P580 (MmpS2), P581/P582 (MmpS4). All clones were sequenced by whole-plasmid sequencing (Plasmidsaurus).

### Electroporation of mycobacteria

Electrocompetent mycobacteria were prepared by 3 washes in 10% glycerol + 0.05% Tween 80. *M. smegmatis* cells were prepared on ice at 4 °C, *M. marinum* and *M. tuberculosis* cells were prepared at room temperature. ∼500 ng of plasmid DNA was added to washed bacterial cell suspensions and incubated for 2 minutes. Electroporation was carried out in a 2 mm gap cuvette with electroporation using a BTX^TM^ ECM 630 electroporation system (Settings: 2.5kV, 1000 Ω, 25 µF). 1 ml prewarmed 7H9 + OADS + Tween 80 was added to the cuvette and incubated at 37°C for 4 hours (*M. smegmatis*), 16 hours (*M. marinum*) and 24 hours (*M. tuberculosis*) before plating on 7H10 + appropriate antibiotics and incubating at 33°C (*M. marinum*) or 37°C (*M. tuberculosis* and *M. smegmatis*). For pFLAG-attP-Kan, pMA-Int (Addgene #110096) was co-electroporated to supply mycobacteriophage L5 integrase for genome integration.

### MmpS5L5 ΔCC expression and lysis

*M. tuberculosis* MmpS5L5 lacking amino acids 494–688 (ΔCC) tagged with a C-terminal GFP-FLAG tag was expressed from an episomal pMEXC3GF vector in *M. smegmatis* mc^2^155. The pMEXC3GF vector was created by replacing the C-terminal His tag from pMEXC3GH (Addgene # 110082, a gift of M. Seeger) with a FLAG tag. 500 ng pMEXC3GF *Mtb* MmpS5L5 ΔCC was electroporated into *M. smegmatis* mc^2^155 as described, and plated on 7H10 agar supplemented with 10% OADS and 50 µg ml^-1^ Hygromycin B. Plates were incubated for 4–5 days at 37 °C. All colonies from the plate were scraped, vortexed to disperse clumps, added to 50 ml 7H9 + 10% OADS + 50 µg ml^-1^ Hygromycin B and incubated for ∼ 24 h at 37 °C. 4 ml pre-culture was added to each of 12 L of 2×TY supplemented with 50 µg ml^-1^ Hygromycin B, 0.1% Tween 80 and 0.01% antifoam 204 (Sigma #A6426). Cultures were incubated at 37 °C with shaking at 170 rpm. At exactly OD_600_ = 0.8, cultures were induced with 1 mg ml^-1^ Anhydrotetracycline hydrochloride (ATc) in DMSO, to a final concentration of 500 ng ml^-1^ ATc. The temperature was reduced to 18 °C and cultures were incubated for 96 h. Bacteria were collected by centrifugation at 8,000 ×g, washed in PBS, and cell pellets stored at −70 °C.

Cell pellets were thawed and resuspended in 5 ml ice-cold lysis buffer per gram of wet cell mass. Lysis buffer: 50 mM HEPES pH 7.4 (4°C), 10% (w/v) sucrose, 2 mM EDTA, SIGMAFAST^TM^ protease inhibitors (Sigma #S8830). Resuspended cells were stirred until homogeneous and filtered through a 425 µm aperture mesh stainless steel sieve (Endecotts Ltd). Cells were lysed by three passes through a Microfluidizer M110-P (Microfluidics) at 25 kPsi, in a 4°C cold-room. All subsequent steps were carried out on ice at 4°C. Unbroken cells and cell wall debris was removed by centrifugation at 8,000 ×g for 40 mins in a Fiberlite F14-6x250Y rotor (Thermo Scientific). Membrane vesicles were collected from the supernatant by ultracentrifugation at 120,000 ×g for 2 hours in a Ti45 rotor (Beckmann Coulter). Membranes were resuspended in 10 ml lysis buffer per gram wet mass of membranes using a Potter-Elvehjem homogeniser (GPE scientific limited, #22010). Protein concentration of the membrane fraction was determined using a bicinchoninic acid (BCA) assay kit (Thermo Scientific #A55865), with BSA standards, according to the manufacturer’s instructions. Membranes were flash-frozen in liquid nitrogen and stored at −70 °C prior to purification.

### MmpS5L5 ΔCC purification

Membranes were thawed and immediately placed on ice. All purification steps were carried out at 4 °C. NaCl was added to a final concentration of 150 mM. 10% Lauryl maltose neopentyl glycol (LMNG) (Anatrace #NG310) was added to a ratio of 1.6 g/g detergent : membrane (∼ 0.3% (w/v) final). Membranes were solubilised for 1 hour at 4 °C on a roller. Insoluble material was pelleted by ultracentrifugation at 120,000 ×g for 30 mins using a Ti70 rotor (Beckman Coulter). Solubilised membranes were incubated with 2 ml anti-FLAG M2 affinity resin (Sigma, #A2220) equilibrated in wash buffer (50 mM HEPES pH 7.4 (4°C), 150 mM NaCl, 2 mM EDTA, 0.002% LMNG) for 2 hours. The resin was collected in a gravity column and rapidly washed with 50 column volumes (CV) of wash buffer. The protein was eluted by addition of 1.5 CV wash buffer supplemented with 200 µg ml^-1^ 3xFLAG peptide (Sigma, #F4799) and incubation for 2 × 30 mins, followed by an overnight elution step. Elution fractions were concentrated using an Amicon Ultra-4 100K MWCO centrifugal concentrator (Millipore, #UFC8100), concentration factor ∼ 24. Aggregated/insoluble material was pelleted by ultracentrifugation at 120,000 ×g for 20 mins in a TLA55 rotor (Beckman Coulter). 250 µl protein at A_280_ = 7.2 was purified by size exclusion chromatography on a Superose 6 Increase 10/300 column (Cytiva) equilibrated in 50 mM HEPES pH 7.4 (4°C), 150 mM NaCl, 0.004% LMNG. Peak fractions were concentrated using a Amicon Ultra-4 100K MWCO centrifugal concentrator to A_280_ = 2.2 and used immediately for cryo-EM grid preparation.

### Cryo-EM sample preparation and data acquisition

For ‘substrate-incubated’ MmpL5, bedaquiline was added to a final concentration of 50 µM, 1.38% dimethylsulfoxide (DMSO). For ‘apo’ MmpL5 this step was omitted. 3.5 µl of MmpL5 at A_280_ = 2.2 were applied to UltrAuFoil 300 mesh gold R1.2/1.3 grids (Quantifoil) that had been freshly glow discharged using an Edwards S150B glow discharger for 70 s at setting 6, 30–35 mA, 1.2 kV and 0.2 mBar (0.15 Torr). The grids were then flash frozen in liquid ethane using a Thermo Fisher Scientific Vitrobot IV maintained at 4°C and 100% humidity (3.5 s blotting time, −15 blotting force) using Whatman filter paper 1.

Data collection was performed using a ThermoFisher Titan Krios G4 transmission electron microscope with a C-FEG operated at 300 kV and equipped with a Falcon 4i direct electron detector and a Selectris X energy filter. Images were collected using the EPU software package in counting mode at a magnification of 130,000× with a physical pixel size of 0.955 Å pixel^-1^, with a total electron dose of 80 electrons per Å^2^ during a total exposure time of 6.20 s, dose-fractionated into 72 movie frames. A slit width of 10 eV on the energy filter and a defocus range of 0.6–2.2 μm with increments of 0.2 µm. A total of 6,071 movies (Bedaquiline dataset) and 6,769 movies (Apo dataset) were collected over two separate sessions.

### Cryo-EM image processing

Data was processed in cryoSPARC v4.6.2 (70). Movies were aligned, drift corrected, and dose-weighted using Patch Motion Correction. Defocus and contrast transfer function (CTF) were estimated using patch CTF estimation. For the Bedaquiline dataset, particles were initially picked using blob picker (particle diameter 110–250 Å) and sorted by 2 sequential rounds of 2D classification. Classes with clear trimeric density were selected (15,100 particles) and used to generate an initial model (1 class). Non uniform refinement produced a 3D volume of a trimer (3.9 Å) that was highly anisotropic (cFAR: 0.01) – and lacked side views. 2D templates were generated from this volume using cryoSPARC’s ‘create templates’ job and used for template picking (templates low-pass filtered to 20 Å), yielding 3,240,490 particles. These particles were subjected to 3 rounds of heterogeneous refinement (6 classes) with the previously generated trimer volume, a monomer volume, and 4 junk ‘decoy’ classes, selecting the best trimer class after each iteration to yield 155,452 particles. Well-defined classes after 2D classification were selected to generate a less anisotropic initial model (1 class). A further 5 rounds of heterogeneous refinement, using the improved initial model with 4 junk ‘decoy’ classes, was performed until < 1% particles were being assigned to decoy classes. The resulting 52,000 particles were further classified by 2D classification, and those classes with well-defined features were selected (35,500 particles), re-extracted with re-centring and used for initial model generation (1 class). Non-uniform refinement produced a 3.3 Å reconstruction that was still anisotropic (cFAR: 0.01), but with improved density features. To preserve more views in the reconstruction, 2D templates were generated using the 3.3 Å trimer volume followed by template picking, yielding 4,100,758 particles. 4 rounds of heterogeneous refinement were performed using the trimer model (low pass filtered to 15 Å), and 7 junk decoy classes, yielding 103,577 particles. A single round of 2D classification was performed and well-defined classes (43,314 particles) were selected, re-extracted with re-centring and used to generate an initial model (1 class). Non-uniform refinement produced a 3.28 Å reconstruction (cFAR: 0.02). A subsequent round of heterogeneous refinement with two initial models (low-pass filtered to 12 Å) resulted in 33,185 (76.8%) particles being assigned to one class. Non-uniform refinement of this class produced a 3.23 Å reconstruction that was less anisotropic (cFAR: 0.04). Reference-based motion correction, followed by non-uniform refinement produced a 3.08 Å map (cFAR: 0.05). For each map, the overall resolution reported in cryoSPARC was estimated using the gold-standard FSC criterion (FSC = 0.143).

For the ‘apo’ dataset, the ‘bedaquiline-trimer’ map was used to generate templates for particle picking yielding 4,810,392 particles. 4 rounds of heterogeneous refinement, using the bedaquiline-trimer map as an initial model, with 7 junk ‘decoy’ classes was performed, yielding 87,473 particles. A subsequent round of 2D classification was performed and well-defined classes were selected (39,277 particles), re-extracted with re-centring and used for initial model generation. Non-uniform refinement yielded a 3.44 Å reconstruction (cFAR: 0.05). Reference-based motion correction followed by non-uniform refinement produced a 3.39 Å reconstruction. A final round of 2D classification and selection of non-junk classes (27,426 particles) followed by non-uniform refinement produced a 3.33 Å reconstruction (cFAR 0.17) with improved density features.

### Model building and refinement

An AlphaFold2 model of a trimer of full-length MmpS5L5 (Model archive accession: ma-l7itj) was used as the starting point for model building. MmpL5 residues 494–688 were removed as they are not present in the construct. Unstructured residues 1–20 of MmpL5 are not visible in the density and were removed. MmpS5 residues 32–142 were removed as they are not resolved in the density. Each subunit was individually rigid-body-fitted into the density in Coot (v.0.9.8.93) (71). Residues 942–952 at the C-terminus of MmpL5 were re-built manually in Coot. The extra density corresponds to MSMEG_4326 (*acpM*), as was identified previously (24). An AlphaFold model of MSMEG_4326 was rigid body fitted into this density in Coot. Phospholipids were identified based on Y shaped densities in the EM map. The length of the acyl chain was modified to fit into where the density for the acyl chain is well resolved. Real-space refinement as performed using PHENIX (v.1.21.2) (72), with Ramachandran and secondary structure restraints enabled, followed by inspection and manual building in Coot. Iterative rounds of real-space refinement and manual building were repeated until satisfactory. Model validation was performed using MolProbity (73). Statistics are summarised in Supplementary Table S3.

### Focused ion beam (FIB-)milling and electron cryotomography (cryo-ET)

To prepare cells for cryo-ET, *M. tuberculosis* mc^2^6206 was grown in 7H9 medium supplemented with OADS without detergent to an optical density of OD_600_=0.5. Clumps of bacteria were removed through centrifugation at 100 g for 10 min, and cells were concentrated through centrifugation at 3,200 g for 10 mins followed by resuspension in fresh medium to a final optical density of OD=3. Cells were then plunge-frozen into liquid ethane using Quantifoil SiO_2_ 200-mesh R 1/4 cryo-EM grids and a Vitrobot Mark IV with a hydrophobic inset on the front-blotting pad to result in back-side blotting. Samples were milled using a ThermoFisher Scientific Aquilos 2 Dual-Beam focused ion beam-scanning electron microscopy (FIB-SEM) system. Tomographic data acquisition on lamellae was carried out using a ThermoFisher Scientific Krios G3 electron microscope equipped with a Gatan BioQuantum energy filter and a Gatan K3 direct electron detector. Tilt series were acquired from −60° to +60° degrees after accounting for the pre-tilt of the lamella using 3° increments and a target dose of 120 e/Å^2^ per tilt series, with a defocus range of −6 to −8 µm. Movies of each tilt image were aligned with MotionCor2 (74). Tilt series alignment and tomogram generation were performed with RELION-5 (75,76). Tomograms were denoised with CryoCare (77). Final tomograms were binned to a pixel size of 12 Å, and boxes along the cell envelope were manually clicked in RELION, followed by extraction of overlapping boxes using the helical extraction options. Two-dimensional alignment and averaging of the extracted cell envelope images led to the final average shown in Figure 2. This image analysis was repeated several times, in different sections of the cell envelope, which yielded approximately the same spacings between each layer of the cell envelope of *Mtb*.

### Cavity detection

Cavities in the MmpL5 model were identified using pyKVfinder (78) implemented in ChimeraX using grid spacing 0.6, inner probe radius of 1.4 Å and an outer probe radius of 4.0 Å.

### Purification of MmpL5 coiled-coil domain (492–683)

*E. coli* codon-optimised N-terminally His-TEV tagged MmpL5-CC (residues 492–683) was expressed in *E. coli* BL21(DE3) as inclusion bodies. Bacteria were grown in 2×TY media supplemented with 100 µg ml^-1^ Ampicillin. Expression was induced at OD_600_ 0.7 by addition of 1 mM isopropyl-β-d-thiogalactoside (IPTG), followed by incubation at 37 °C for 3 hours. Bacteria were collected by centrifugation at 5,000 ×g. Bacterial pellets were suspended in 100 ml lysis buffer (50 mM Tris pH 7.6, 150 mM NaCl, 10 mM EDTA, 10 mM DTT, 1% (v/v) Triton X-100, 0.2 mM PMSF supplemented with a spatula tip of Lysozyme, and 500 units Benzonase nuclease (Sigma). Bacteria were lysed by 2 passes through a microfluidizer M110-P at 25 kPsi. Inclusion bodies were pelleted by centrifugation at 20,000×g and washed by resuspension with 80 ml lysis buffer using a Potter-Elvehjem homogeniser (GPE scientific limited, #22010). Inclusion bodies were pelleted by centrifugation at 20,000 ×g, and stored at −20 °C. Inclusion body was solubilised with 20 ml 50 mM Tris pH 7.6, 150 mM NaCl, 6 M Urea, 10 mM DTT and incubated for 1 hour at room temperature. Insoluble material was removed by centrifugation at 20,000 ×g. DTT was removed by buffer exchange using a Zeba 7,000 MWCO desalting column equilibrated in unfolding buffer (50 mM Tris pH 7.6, 150 mM NaCl, 6 M Urea). Unfolded protein was diluted to a protein concentration of 1 mg ml^-1^ in unfolding buffer before refolding by slow dropwise addition to 3 L refolding buffer (50 mM Tris pH7.6, 150 mM NaCl, 1 mM GSSH, 1 mM GSH) with gentle stirring. Refolding was allowed to proceed for 30 minutes at room temperature. Refolded protein was purified by ion-exchange chromatography, using a HiPrep Q 16/10 column pre-equilibrated in IEX A buffer (50 mM Tris pH 7.6, 150 mM NaCl) and then eluted with a linear gradient to 1M NaCl. Peak fractions were concentrated and dialysed overnight against 5 L storage buffer (50 mM Tris pH 8.0, 150 mM NaCl) with a 3 kDa MWCO dialysis cassette. Protein aliquots were flash frozen and stored at −70°C until use.

### Circular dichroism spectroscopy

Circular dichroism data were collected using a JASCO J-815 spectropolarimeter in the far UV region. Spectra Manager (1.55) was used for data collection. Circular dichroism spectra were acquired at 14 μM protein concentration in 50 mM Tris pH 7.6, 150 mM NaCl at room temperature. Data were collected in a 1-mm quartz cuvette between wavelengths of 190 nm and 260 nm with the instrument set as follows: band width, 1 nm; scanning speed, 50 nm min^−1^; response time, 1 s. Each circular dichroism spectrum was obtained by averaging eight scans and subtracting the background signal of the buffer and cuvette. The spectra were converted from ellipticities (mdeg) to mean residue ellipticities (deg·cm^2^·dmol^−1^·res^−1^) by normalizing for concentration of peptide bonds and the cell path length using the equation:

MRE = θ × 10^6^ / c × l × n, where the variable θ is the measured difference in absorbed circularly polarized light in millidegrees, c is the concentration of the protein in micromolar, l is the path length of the cuvette in millimetres, and n is the number of amide bonds in the polypeptide.

### Intact mass spectrometry

Purified N-terminally His-TEV tagged MmpL5 coiled-coil domain was analysed by mass spectrometry both before and after incubation at 45 °C for 30 minutes in 0.05 mM TCEP. The protein was analysed by LC-MS using a Vanquish HPLC (Thermoscientific, USA) with mobile phase A – 90% water, 9.9% acetonitrile with 0.1% formic acid and mobile phase B – 80% acetonitrile, 19.9% water, and 0.1% formic acid. Delivered flow was 300 μl min^-1^. Proteins were trapped on a MAbPac RP 2.1 x10 mm trapping column for two minutes before separation using a MAbPac RP 2.1 x 50 mm LC column. The gradient was from 10% to 30% B over 2 minutes before increasing to 55% B over the next 8 minutes at 80 °C. Proteins were eluted and ionized with a HESI ion source onto an orbitrap mass spectrometer (Exploris MX, Thermoscientific USA). MS data were acquired in in a broadband detection mode (R = 120,000, m/z 600 – 2000, Intact protein mode) in positive ion mode (AGC = 100%, μscans = 10, in source energy = 10 eV). Protein deconvolution was carried out using a sliding window Xtract (isotopically resolved) deconvolution to generate monoisotopic protein masses.

### Surface plasmon resonance

A Biacore T200 instrument (Cytiva) was used for SPR analysis. All steps were carried out at room temperature. GFP-tagged MmpL5 ΔCC was immobilised on a Sensor Chip SA (Cytiva #BR100531) using a biotinylated anti-GFP nanobody for capture. Biotinylated anti-GFP nanobodies purified and biotinylated as described (79). Nanobody was diluted in running buffer (50 mM HEPES pH 7.5 (at 22 °C), 150 mM NaCl, 0.004 % (w/v) LMNG, 5 % DMSO) to 26 μM and immobilised on the SA chip in both measurement and reference channels with a flow of 30 μl min^-1^ for 10 minutes resulting in an immobilisation level of 5995 and 6170 RU, respectively. GFP was diluted in running buffer to 1.8 μM and captured by the GFP nanobody on the reference channel with a flow of 10 μl min^-1^ for 5 minutes resulting in an immobilisation level of 3821 RU. *Mtb* MmpL5 ΔCC-GFP at 50 μg ml^-1^ was captured by the GFP nanobody on the measurement channel with a flow of 10 μl min^-1^ for 5 minutes resulting in an immobilisation level of 5240 RU. Bedaquiline at 2 mM in DMSO was diluted to 100 μM with 50 mM HEPES pH7.5 (at 22°C), 150 mM NaCl, 0.004% (w/v) LMNG, resulting in 5% DMSO final concentration. This 100 μM stock was diluted in a 2-fold dilution series to 12.5 nM in running buffer containing 5% DMSO. The dilution series were sequentially injected onto the chip at 30 μl min^-1^ for 120 s in the association phase, with a 480 second dissociation time. Each dilution series was measured in triplicate. Data analysis was performed in the Biacore T200 evaluation software (v. 1.0) . DMSO solvent correction was applied (80), and the data were double referenced (81) by subtracting the signal from the reference channel, as well as the buffer only injection. The equilibrium response (R_eq_) was plotted against analyte concentration and fitted with non-linear regression using a specific binding with Hill slope model to determine steady-state binding affinity (K_d_). Curve fitting was performed in GraphPad Prism (v. 10.0.2) using a specific binding with Hill slope model, as assuming a Hill coefficient of 1 produced a poor fit.

### Minimum inhibitory concentration (MIC) assay

Minimum inhibitory concentrations were determined by broth microdilution (82,83). MIC plates were prepared with 2 x concentration of test compound in 7H9 + 10% (v/v) OADS without Tween 80 in polystyrene 96-well plates (Corning, Falcon #351177) as polypropylene plates alter the measured MIC of bedaquiline (84). Plates were sealed and stored at −20 °C. Post-thaw, plates were centrifuged at 500×g for 10 minutes before removal of plate seal. Test strains were diluted to 0.5 McFarland (McF) units in 7H9 + OADS (with 0.05% Tween 80) using a nephelometer. 100 µl 0.5 McF cell suspension was diluted 100-fold in 7H9 + OADS (with 0.05% Tween 80). This 0.5 McF unit cell suspension was diluted with 7H9 + OADS + 48 µg/ml pantothenic acid + 100 µg ml^-1^ L-leucine. 100 µl of this diluted suspension, corresponding to ∼ 5 × 10^5^ CFU, was added to each well of the 96-well plate to obtain a 1× concentration of test compound. Plates were sealed and incubated at 33 °C (*M. marinum*) or 37 °C (*M. tuberculosis*) for 10 days (*M. marinum*) or 14 days (*M. tuberculosis*). Plates were sealed and imaged using an Epson perfection V850 photo scanner. Minimum inhibitory concentrations were determined as the minimum concentration of test compound that results in no growth visible by eye.

### Isolation of intragenic suppressors

Suppressor mutants of N444K were isolated by plating *M. marinum mmpL5*::Tn + pFLAG-*Mtb MmpS5L5* N444K on 25 ng ml^-1^ bedaquiline. ∼3 × 10^9^ CFU were plated and incubated for 25 days at 33 °C. This produced 12 resistant colonies, corresponding to a resistance frequency of 2.3 × 10^-8^. Resistant colonies were validated by re-streaking colonies on 7H10 + OADS + 25 ng ml^-1^ bedaquiline plates together with the parental (N444K) and wild-type MmpS5L5 strains. The integrated *Mtb mmpS5L5* and *atpE* loci were PCR amplified from crude genomic DNA using primers P677/P678 (*mmpS5L5* locus) and P812/P813 (*atpE*). PCR products were purified using a PCR purification kit (Qiagen) and either linear amplicon sequenced (Plasmidsaurus) or Sanger sequenced using M13F primers.

### Error-prone PCR library construction and drug selections

Error-prone PCR was conducted using the Genemorph II random mutagenesis kit (Agilent #200550). 1 µg target sequence with 14 PCR cycles was used to produce an average of 1–4 nucleotide changes per gene. Template plasmid was digested with an excess of DpnI (NEB #R0176) for 2 hours. PCR product was resolved on a 1 % agarose gel and purified by gel extraction with a Monarch DNA gel extraction kit (NEB #T1020) according to manufacturer’s instructions. The resulting purified error-prone amplicon with overhangs compatible with FX cloning (69) was cloned into pMINTF3 with a golden-gate assembly protocol. The golden-gate reaction was transformed into 10 aliquots of chemically competent E. coli DH5-α cells prepared by the method of Inoue (85) (∼ 2 × 10^8^ CFU µg^-1^ DNA) and plated on TYE agar + 50 µg ml^-1^ kanamycin in large bioassay plates. This produced ∼120,000 CFU, which, assuming only 1 nucleotide change per gene, corresponds to ∼9-10× coverage at each nucleotide in the gene. Plasmid DNA from 10–15 colonies was isolated and sequenced using whole-plasmid sequencing (Plasmidsaurus) to estimate mutation frequency. All colonies were scraped from the surface of the plate and plasmid DNA for the library was isolated.

∼1 µg plasmid library DNA was electroporated into *M. marinum mmpL5*::Tn cells in 10 replicate electroporations and recovered for 16 hours at 33°C with the addition of 1 ml 7H9 media. The recovered cells were pooled and plated directly on 7H10 agar supplemented with 200 ng ml^-1^ bedaquiline in large bioassay plates to select for resistant bacteria. The library size in *M. marinum* was calculated by CFU enumeration on 7H10 agar + 25 µg ml^-1^ kanamycin which resulted in 7.6 × 10^4^ Kan^R^ colonies. Plates were incubated for 30 days at 33°C. Bedaquiline resistant colonies were inoculated into 7H9 media + 25 µg ml^-1^ kanamycin and sequencing of the integrated complementation vector was performed. Analysis of sequence differences was performed using Snapgene’s “align to reference sequence” tool.

### Immunoblot analysis

Protein concentration was measured using bicinchoninic acid (BCA) assay (Thermo-Scientific) kit according to manufacturer’s instructions, using bovine serum albumin standards (Thermo-Scientific #23208) as a calibration curve. Protein was resolved by SDS-PAGE before being transferred to 0.2 µm PVDF membranes (BioRad #1704156) by semi-dry blotting using a Transblot turbo transfer system (BioRad). Membranes were blocked using 3% (w/v) skimmed milk powder (Sigma #70166) in PBS. Membranes were incubated with primary antibodies at appropriate dilutions in 3% (w/v) skimmed milk powder in PBS for 1 hour at 22°C or overnight at 4°C. Membranes were washed using PBS supplemented with 0.1% (v/v) Tween 20. Membranes were incubated with secondary antibodies at appropriate dilutions in 3% (w/v) skimmed milk powder in PBS for 1 hour at 22 °C or overnight at 4°C and washed with PBS supplemented with 0.1% (v/v) Tween 20. Blots were imaged at 700 and 800 nm using a Licor Odyssey imaging system. 3xFLAG tags on MmpL5 were detected using a monoclonal mouse anti-FLAG M2 antibody (Sigma-Aldrich #F1804) at 1 μg/ml (1:1000). *Mtb* GroEL2 was detected using the IT-56 (CBA1) mouse monoclonal antibody at 1:1,000 dilution (BEI Resources, NIAID, NIH: NR-13655). Goat anti-mouse IgG conjugated with Dylight™ 800 4x PEG (Cell signalling technology #5257) was used at 0.1 μg/ml (1:10,000).

### Sequence alignment and analysis

Protein sequences were aligned using Clustal Omega (86) and analysed with Jalview (87).

### Protein structure prediction

Structure prediction of MmpL5 and MmpS5 complexes was performed using a local implementation of Colabfold version 1.5.2 (88) using MMseqs2 (89) for multiple sequence alignment generation and AlphaFold2 multimer version 3 for prediction of protein complex structures (90). PAE plots were generated using the PAE Viewer webserver (91).

### Data, Materials, and Software Availability

All unique/stable reagents generated in this study are available without restriction from the corresponding authors. Genbank files for the plasmid DNA constructs listed in table S1 is available at: https://doi.org/10.5281/zenodo.15720311. The cryo-EM maps have been deposited at the Electron Microscopy Data Bank under the following accession codes: EMD-53941 (Trimer, apo), EMD-53947 (Trimer, bedaquiline-incubated). The coordinates of the atomic models have been deposited at the PDB under the following accession codes: 9RFU (Trimer, apo), 9RGB (Trimer, bedaquiline-incubated). The AlphaFold2 model of the MmpS5L5 trimer is available in ModelArchive (92) at https://www.modelarchive.org/doi/10.5452/ma-l7itj.

## Acknowledgments and funding sources

We would like to thank Chris Tate for advice on membrane protein purification. We are grateful to Brendan Cormack, Morwan Osman, Alex Lake and Jonathan Shanahan for advice and encouragement. A.J.F thanks the LMB media kitchen for their invaluable help preparing media and plates. We acknowledge the LMB electron microscopy facility for sample preparation and data collection. We thank Toby Darling, Jake Grimmett, and Ivan Clayson (LMB scientific computing) for computing support. We thank Matt McNeil for the gift of TBAJ-876.

B.F.L was supported by Wellcome Trust Investigator Awards (222451/Z/21/Z) and an ERC Advanced Award (742210). L.R was supported by a Wellcome Trust Principal Research Fellowship (223103/Z/21/Z). Research reported in this publication was supported by the National Institute of Allergy and Infectious Diseases of the National Institutes of Health under Award Number R01AI054503 (L.R). The content is solely the responsibility of the authors and does not necessarily represent the official views of the National Institutes of Health. This work was supported by the Medical Research Council, as part of United Kingdom Research and Innovation (also known as UK Research and Innovation) [Programme MC_UP_1201/31 to T.A.M.B.]. T.A.M.B. would like to thank the European Molecular Biology Organization, the Wellcome Trust (Grant 225317/Z/22/Z), the Leverhulme Trust, and the Lister Institute for Preventative Medicine for support. For open access, the author has applied a CC BY public copyright licence to any Author Accepted Manuscript version arising from this submission. This work is licensed under a Creative Commons Attribution 4.0 International Licence.

