## Supplementary material for "Structural and functional analysis of the *Mycobacterium tuberculosis* MmpS5L5 efflux pump presages a pathway to increased bedaquiline resistance": SI Appendix

Lalita Ramakrishnan

Ben Luisi

**This PDF file includes:**

Figures S1 to S19

Tables S1 to S3

SI References

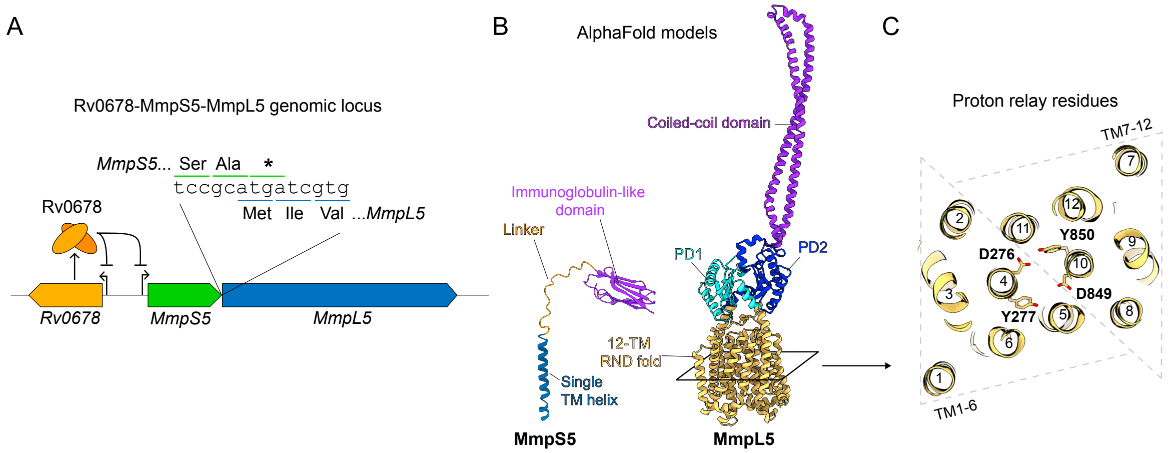

D

| Strain | Bedaquiline MIC (ng ml$^{-1}$) | Clofazimine MIC (ng ml$^{-1}$) | PBTZ-169 MIC (ng ml$^{-1}$) | TBAJ-876 MIC (ng ml$^{-1}$) | TBAJ-587 MIC (ng ml$^{-1}$) | OPC-167832 MIC (ng ml$^{-1}$) | A23187 MIC (µg ml$^{-1}$) | Nigericin MIC (µg ml$^{-1}$) |
|---|---|---|---|---|---|---|---|---|
| Number of replicates | 3 | 3 | 3 | 2* | 3 | 4 | 4 | 4 |
| mc$^2$6206 | 25.0 | 7.8 | 0.78 | 12.5 | 6.25 | 0.19 | 2.0 | 1.0 |
| mc$^2$6206 Δ*S5L5* | 6.25 | 1.95 | 0.39 | 3.12 | 1.56 | 0.19 | 0.5 | 0.5 |
| mc$^2$6206 Δ*S5L5* + Mtb *MmpS5L5* | 50.0 | 7.8 | 1.56 | 12.5 | 6.25 | 0.79 | 8.0 | 2.0 |
| mc$^2$6206 Δ*S5L5* + Mtb *MmpS5L5* Y277F | 50.0 | 7.8 | 0.39 | 12.5 | 6.25 | N.D | N.D | N.D |
| mc$^2$6206 Δ*S5L5* + Mtb *MmpS5L5* D849N | 6.25 | 1.95 | 0.39 | 3.12 | 1.56 | N.D | N.D | N.D |
| mc$^2$6206 Δ*S5L5* + Mtb *MmpS5L5* Y850F | 6.25 | 1.95 | 0.39 | 3.12 | 1.56 | N.D | N.D | N.D |

* - Not in triplicate due to limited reagent

**Fig. S1: MmpS5L5 is a multidrug efflux pump**

(A) Diagram of the Rv0678 MmpS5 MmpL5 genomic locus. MmpS5 and MmpL5 are translationally coupled with overlapping stop-start codons. (B) AlphaFold models of *M. tuberculosis* MmpS5 (Uniprot: P9WJS7) and MmpL5 (Uniprot: P9WJV1) with structural features annotated. (C) View of slice through the transmembrane domain of MmpL5, with TM helices numbered. In TMs 4 and 10, Asp-Tyr 'Proton-relay' residues are shown. (D) MIC values for knockout, complemented strains performed in biological triplicate. MICs are given in either ng ml$^{-1}$ or µg ml$^{-1}$.

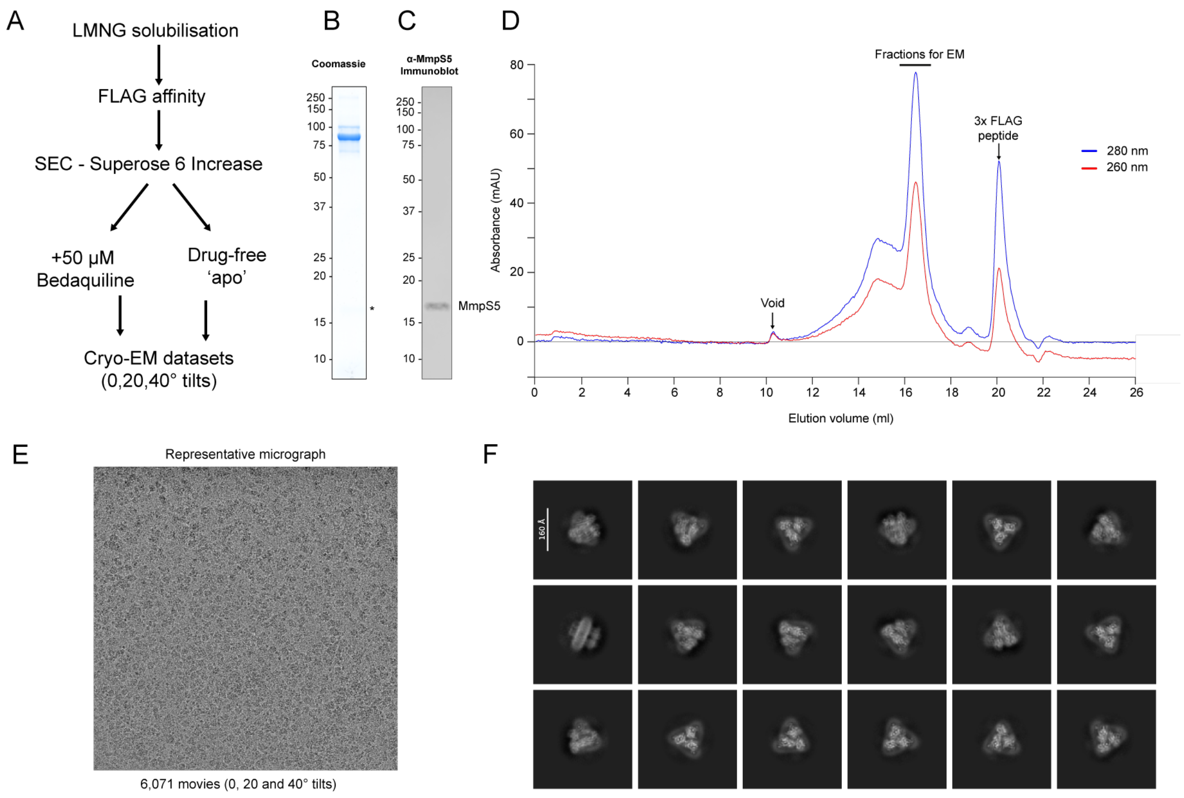

**Fig. S2: Purification of MmpS5L5 ∆CC**

(A) Purification scheme for MmpS5L5-GFP-FLAG. (B) Coomassie-stained SDS-PAGE peak fraction used for cryo-EM sample preparation. (C) Anti-MmpS5 immunoblot of fractions used for cryo-EM sample preparation. (D) Size-exclusion chromatography profile of affinity purified MmpL5-GFP-FLAG on a Superose 6 10/300 Increase column. (E) Representative cryo-EM micrograph of the +bedaquiline dataset. (F) 2D class averages for the trimeric species of MmpL5 in the dataset.

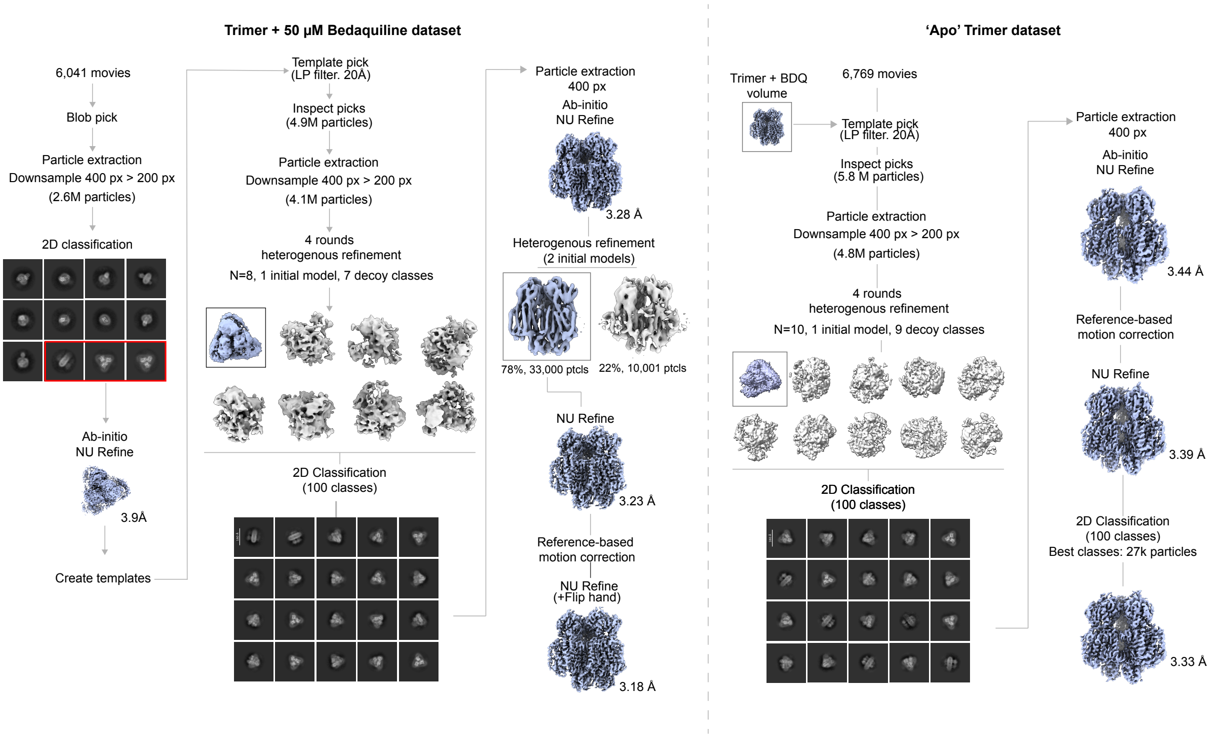

### Fig. S3: Cryo-EM processing pipeline

Cryo-EM processing workflow (cryoSPARC) for both apo and bedaquiline-incubated trimer datasets.

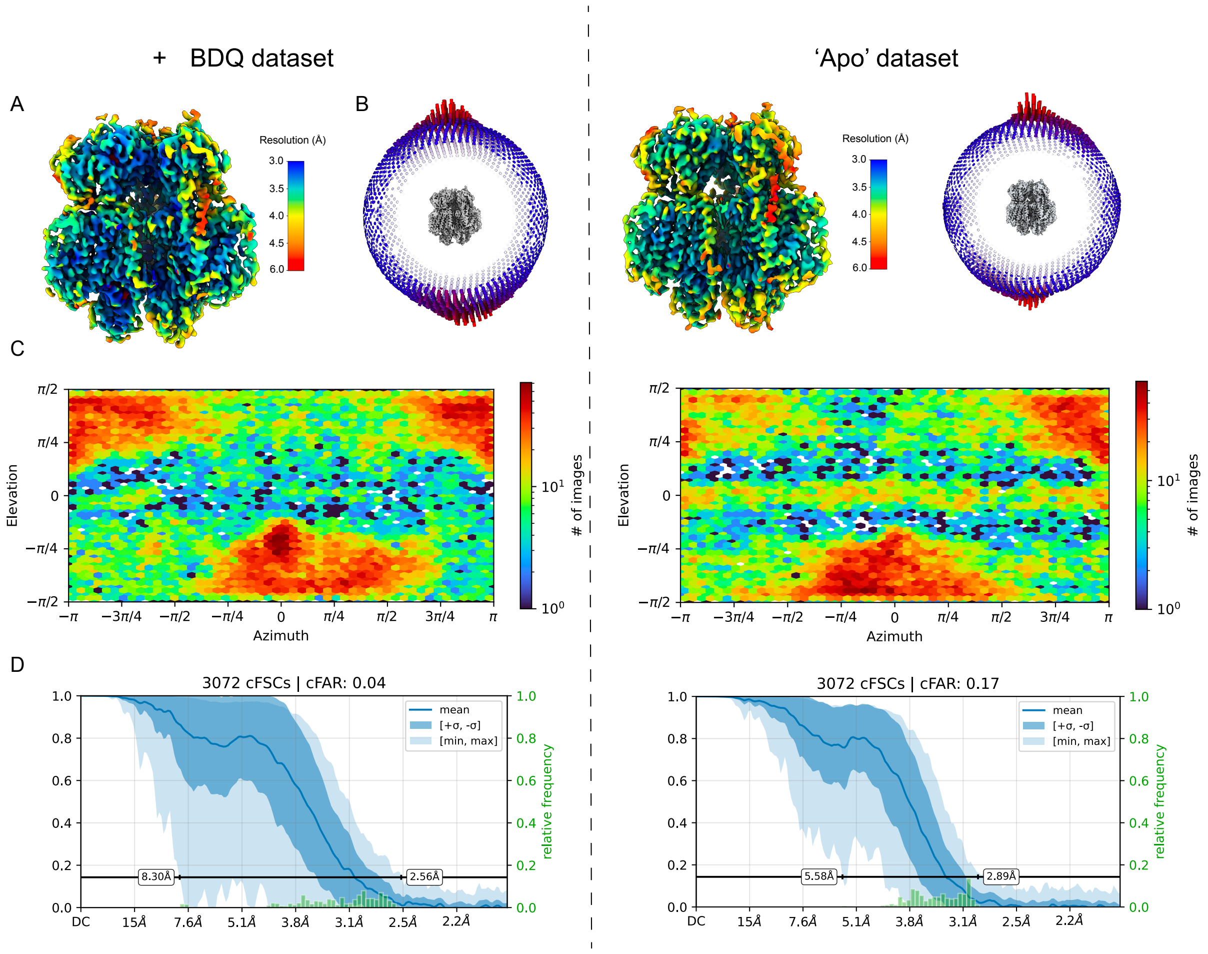

**Fig. S4: Local resolution, angular distributions, and FSC curves.**

(A) Cryo-EM density map coloured according to local resolution. TM helix 2 and PD2 have lower resolution, consistent with flexibility in these regions. (B–C) Angular distributions of the final reconstruction. (D) Fourier shell correlation (FSC) plot of the trimer reconstruction.

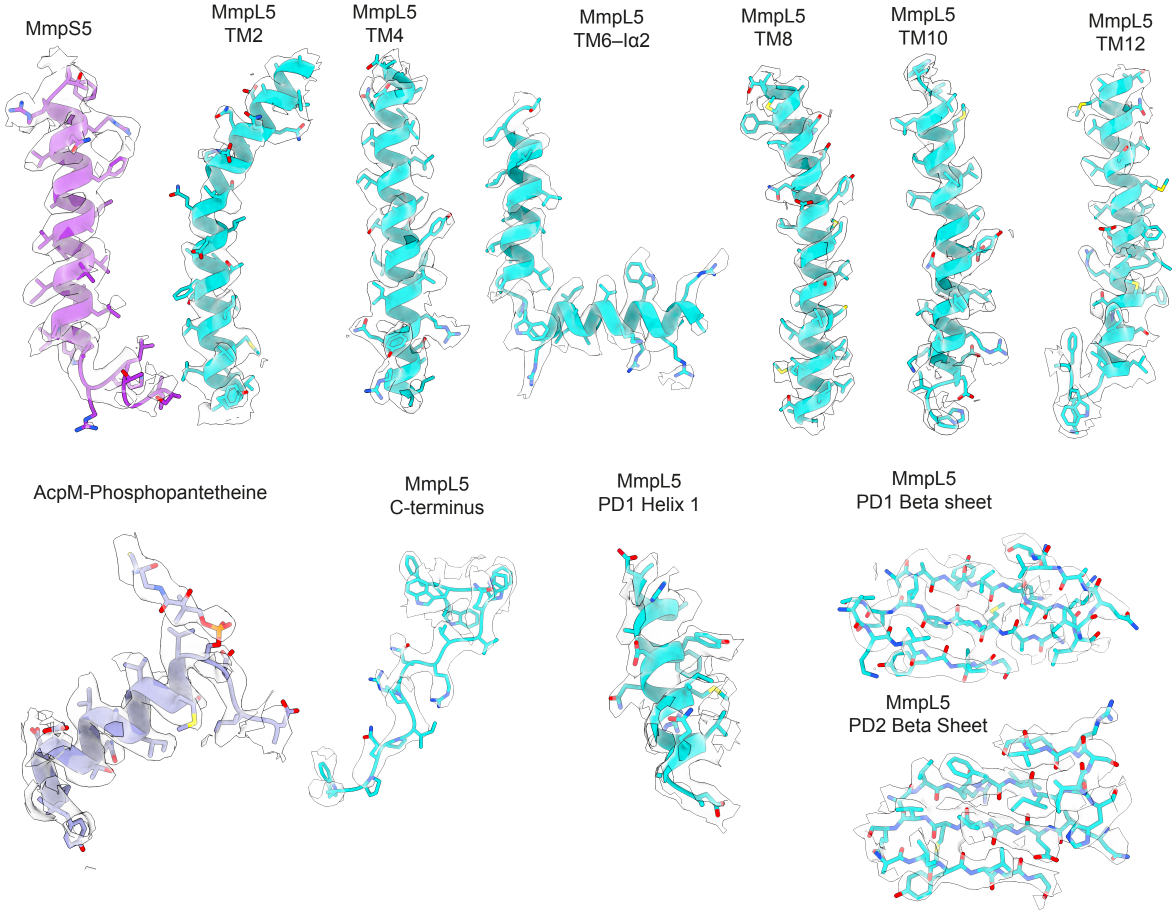

**Fig. S5: Model-map fit for MmpS5, MmpL5 and AcpM**

Representative map-model fit for cryo-EM densities from the bedaquiline-incubated trimer map. Density was extracted using ChimeraX's 'volume zone' tool with a 2.0–3.0 Å distance cutoff around the indicated secondary structure.

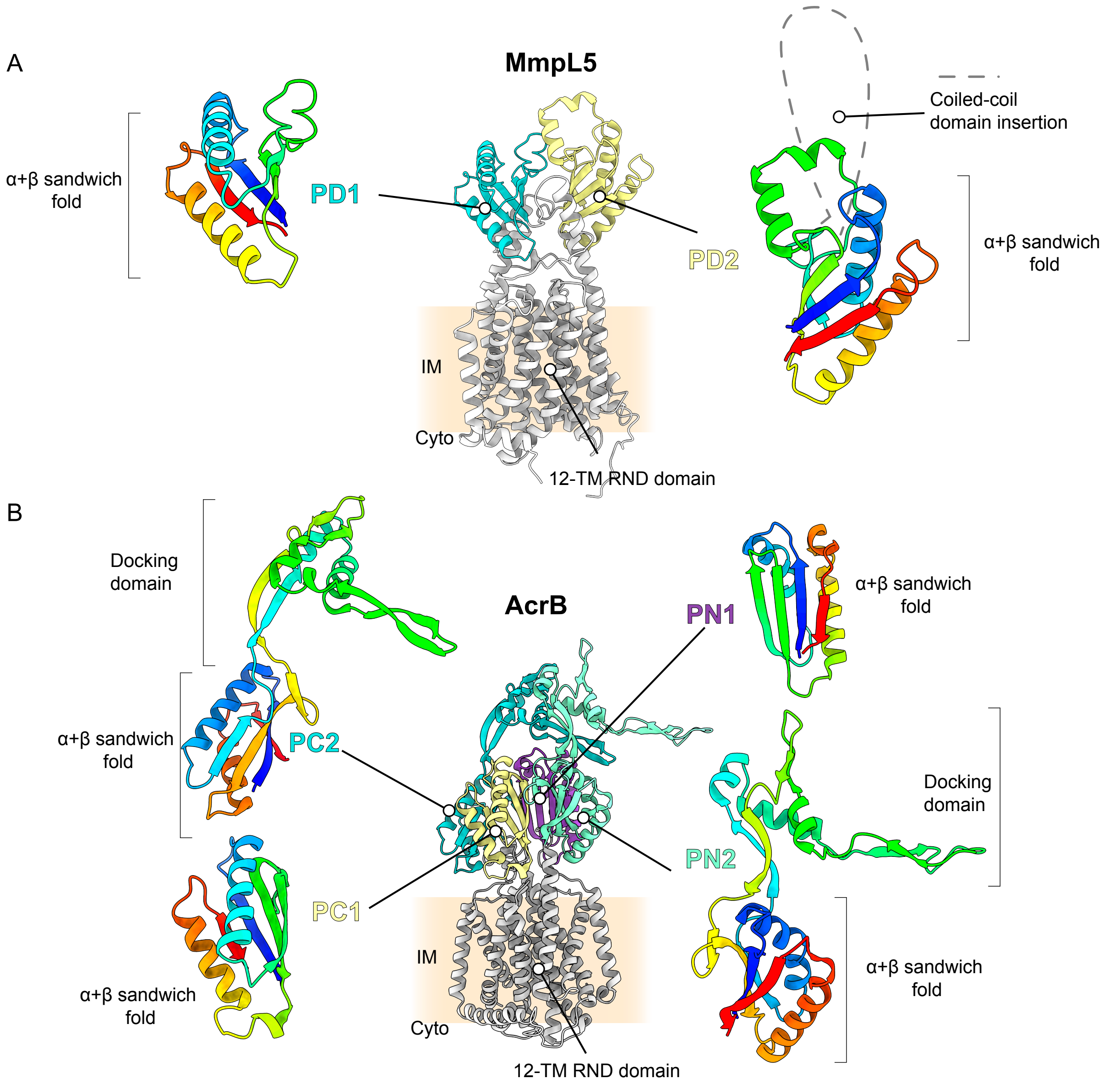

**Fig. S6: Comparison of AcrB and MmpL5**

Upper, Domain structure of *Mtb* MmpL5 ΔCC. The coiled-coil domain inserted into PD2 is shown as a dashed line. Lower, Domain structure of an *E. coli* AcrB protomer. Individual domains are coloured with a rainbow gradient from blue at the N-terminus to red at the C-terminus.

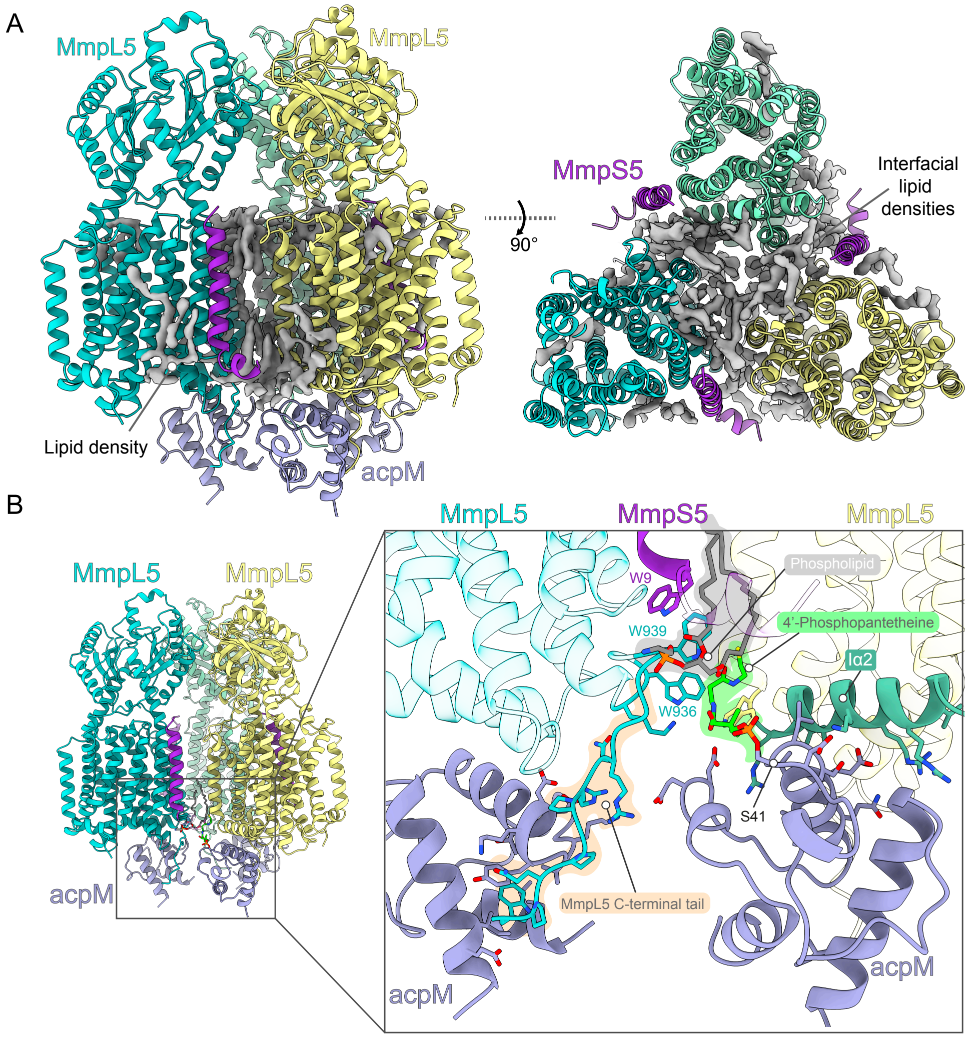

**Fig. S7: Lipids and AcpM stabilise intersubunit interactions**

(A) Structural model of MmpS5L5-AcpM with unassigned lipid-like densities (grey) at the interfaces between subunits. (B) Interface between protomers at the inner leaflet of plasma membrane. AcpM Ser41 is post-translationally modified with 4'-Phosphopantetheine (green). A phospholipid (grey) interacts with Trp9 of MmpS5 (purple). AcpM (lilac) interacts with MmpL5's intracellular Iα2 helix (teal), and MmpL5's C-terminal tail (blue/orange).

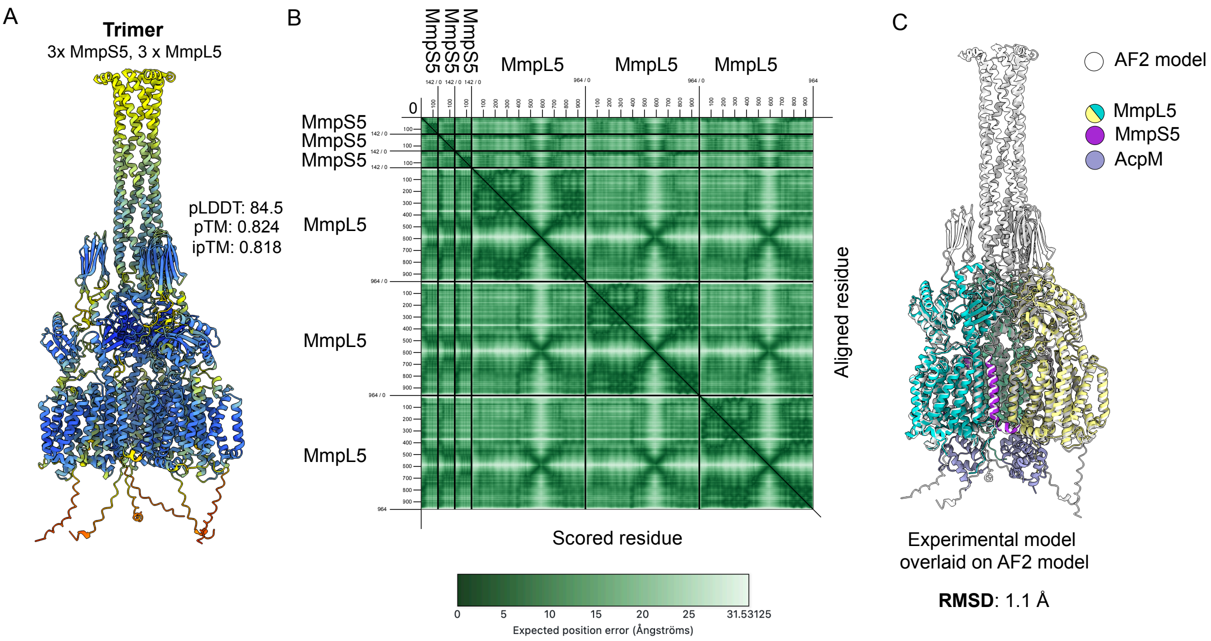

**Fig. S8: Alphafold2 prediction of a homotrimer of MmpS5 and MmpL5**

(A) Structural model of a MmpS5L5 trimer. Models are coloured by per-residue pLDDT score. The pTM and ipTM values for the corresponding model are indicated. (B) Predicted aligned error (PAE) plot of the trimeric model. (C) Overlay of the experimentally determined structure of MmpS5L5-AcpM, on the AlphaFold2 model. RMSD – Root mean square deviation.

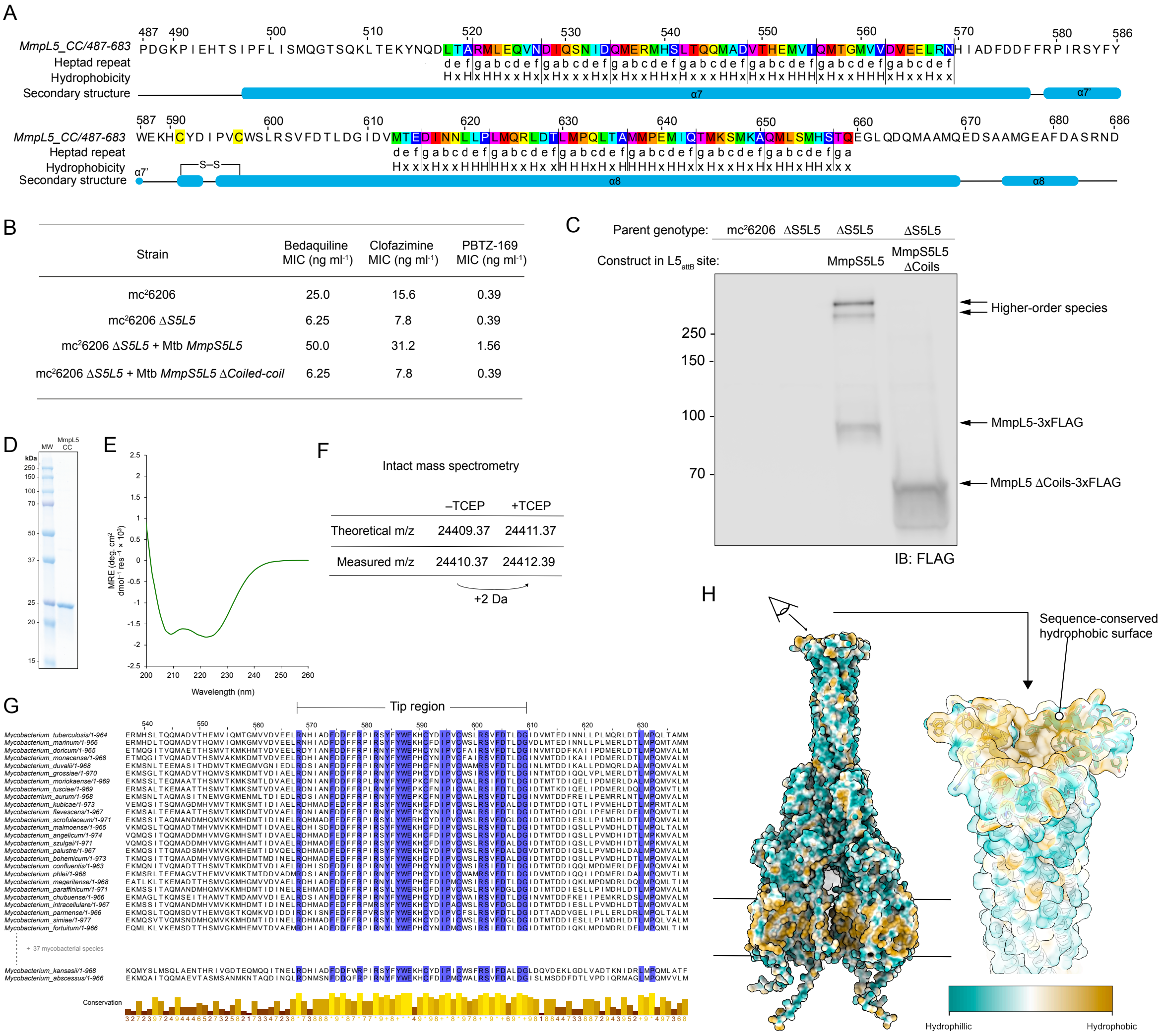

**Fig. S9: Coiled-coil heptad repeat sequence analysis and functional validation.**

(A) Annotated amino acid sequence of *M. tuberculosis* MmpL5's coiled-coil domain (residues 487 – 683). Hydrophobic amino acids are indicated with "H". Non-hydrophobic amino acids are indicated with 'x'. Heptad repeats are classified according to (1). gabcef indicates the position within the heptad repeat. S–S disulfide bond. (B) Bedaquiline, clofazimine and PBTZ-169 MIC values for strains lacking residues (494–687) (ΔCC). (C) anti-FLAG immunoblot showing that the ΔCC construct is expressed. (D) Coomassie stained SDS-PAGE gel of purified MmpL5 coiled-coil domain. (E) CD spectrum of MmpL5 coiled-coil domain. (F) Theoretical and measured monoisotopic masses of MmpL5 coiled-coil domain, with and without TCEP reduction. (G) Multiple sequence alignment of MmpL5 from a wide sampling of mycobacterial species. Residues 537-638 are shown. 100% sequence conserved amino acids are coloured blue. Conservation score is shown below. (H) Surface of the MmpS5L5 Alphafold2 model coloured according to the Kyte-Doolittle hydrophobicity scale (2).

A

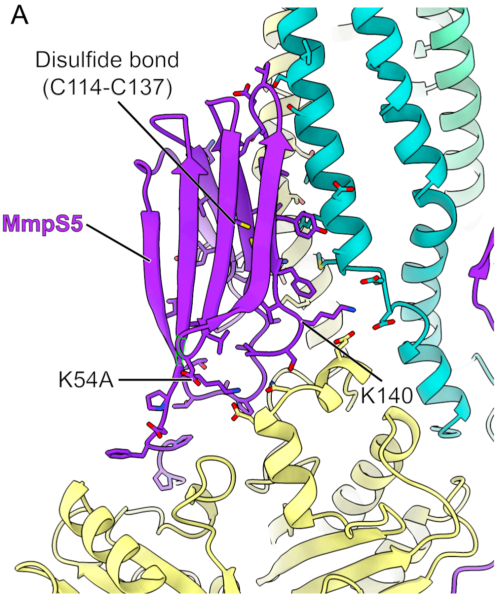

B

| Strain | Bedaquiline MIC (ng ml$^{-1}$) | Clofazimine MIC (ng ml$^{-1}$) | PBTZ-169 MIC (ng ml$^{-1}$) |
|---|---|---|---|
| mc$^2$6206 | 25.0 | 15.6 | 0.39 |
| mc$^2$6206 Δ*S5L5* | 3.1 | 7.8 | 0.39 |
| mc$^2$6206 Δ*S5L5* + Mtb *MmpS5L5* | 50.0 | 31.2 | 1.56 |
| mc$^2$6206 Δ*S5L5* + Mtb *MmpL5* only | 3.1 | 7.8 | 0.39 |
| mc$^2$6206 Δ*S5L5* + Mtb *MmpS5* K54A *MmpL5* | 6.3 | 7.8 | 0.39 |
| mc$^2$6206 Δ*S5L5* + Mtb *MmpS5* K140A *MmpL5* | 25.0 | 7.8 | 0.79 |
| mc$^2$6206 Δ*S5L5* + Mtb *MmpS5 K54A K140A MmpL5* | 3.1 | 7.8 | 0.39 |
| mc$^2$6206 Δ*S5L5* + Mtb *MmpS5 C137S MmpL5* | 3.1 | 7.8 | 0.39 |

C

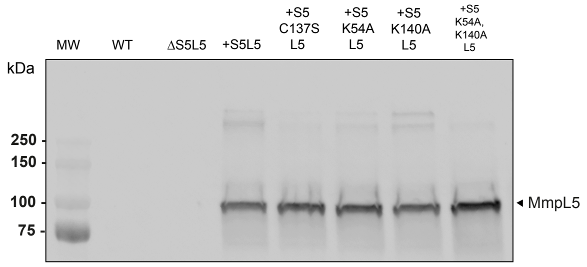

**Fig. S10. Mutations at the S5-L5 interface prevent complementation**

(A) Structural model of the MmpS5-MmpL5 interface from the AlphaFold2 prediction. (B) Bedaquiline, clofazimine and PBTZ-169 MIC values for selected MmpS5 mutations. (C) Anti-FLAG immunoblot showing MmpL5 expression in all strains.

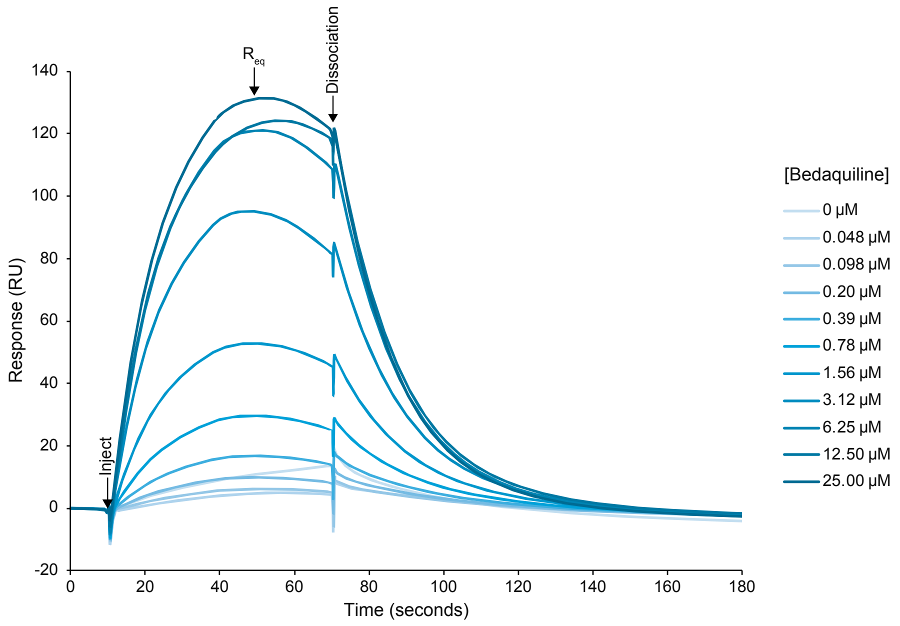

**Fig. S11. Representative SPR sensorgram of bedaquiline binding to *Mtb* MmpL5 ΔCC-GFP**

Sensorgram from a representative surface plasmon resonance (SPR) experiment, showing the binding of bedaquiline to an *Mtb* MmpL5 ΔCC-GFP. The x-axis represents time in seconds, and the y-axis represents the response in resonance units (RU). Different concentrations of bedaquiline, ranging from 0.024 μM to 25.00 μM, were injected over the sensor chip, as indicated by the legend on the right. We have observed that the response begins to decrease before the dissociation phase in multiple independent experiments. The peak of the response is used as the $R_{eq}$ value.

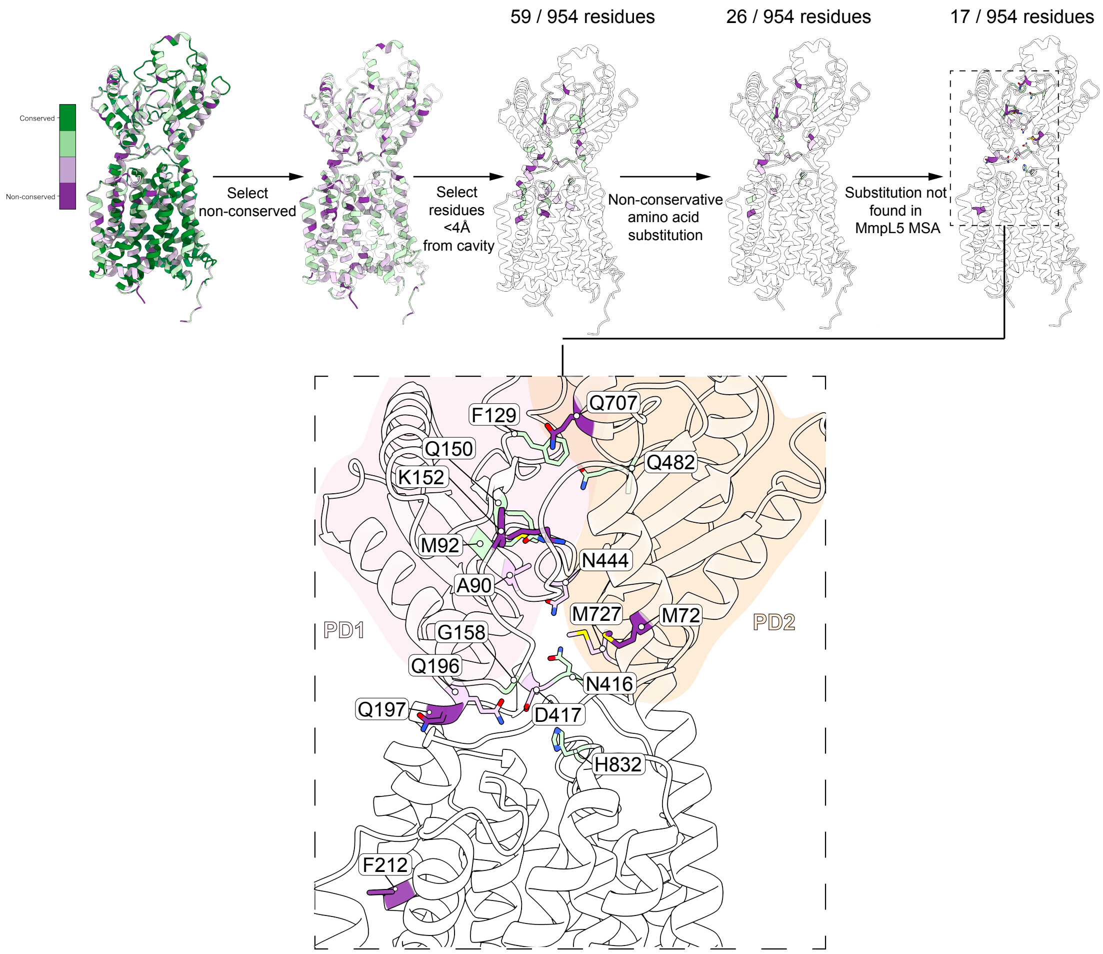

**Fig. S12. Paralog-guided mutation selection criteria**

(A) Selection criteria for identifying candiate residues with functional importance. Sequence conservation between MmpL5/4/2/1 was mapped onto the structure of MmpL5. 100% sequence conserved residues were discounted. Residues within 4 Å of the cavity formed by PD1, PD2 and TMs 1–4 (orange) were selected. Of these, those with non-conservative amino acid substitutions within an MSA of MmpL5/4/2/1 were selected. Selected substitutions that were found in MmpL5s from other mycobacterial species were discounted. (B) Structural model of MmpL5, showing the positions of selected residues based on the selection criteria.

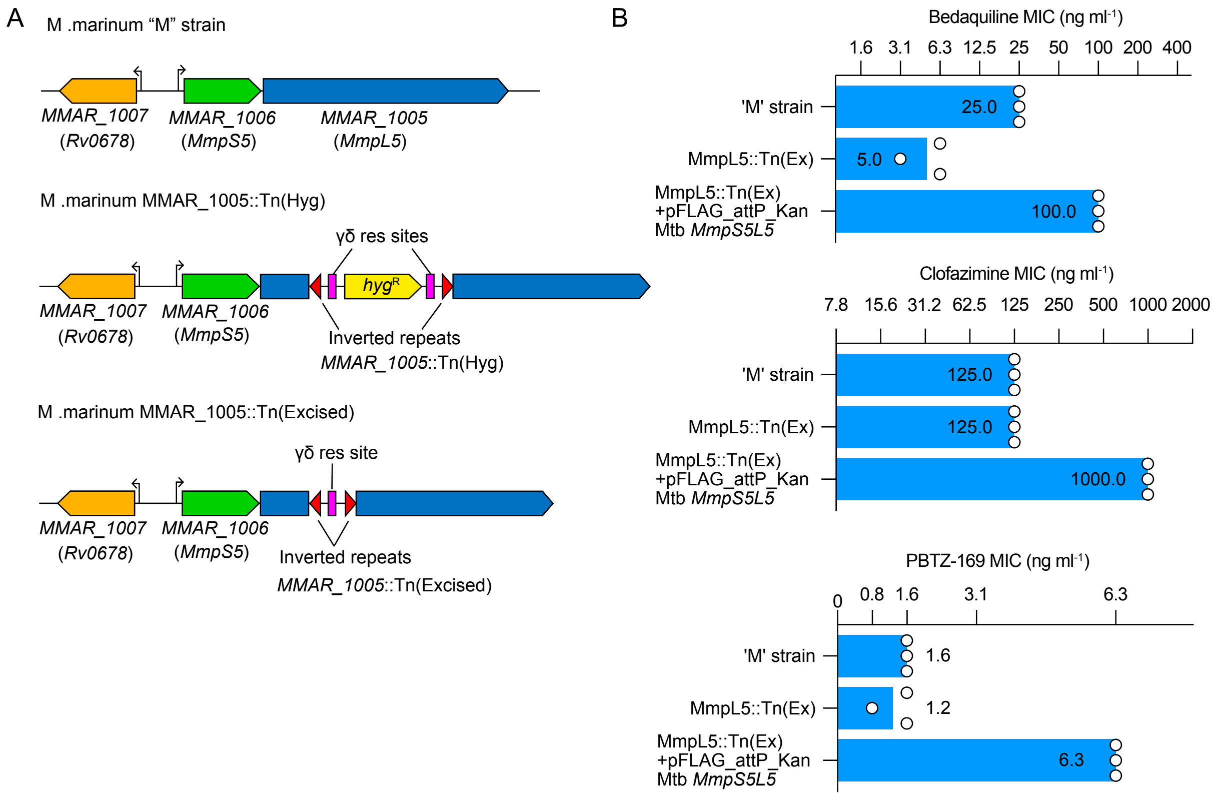

**Fig. S13. *M. marinum* is a suitable surrogate for MmpS5L5 phenotypes**

(A) Schematic of each *M. marinum* transposon strain used in this study. (B) Bar graphs of MIC values for *M. marinum* strains complemented with pFLAG-attP-Kan *Mtb MmpS5L5* integrated at the L5$_{attB}$ site for bedaquiline, clofazimine and PBTZ-169. Open circles are plotted MIC values for independent repeats. Values for each column indicate the geometric mean of the MIC values in ng ml$^{-1}$.

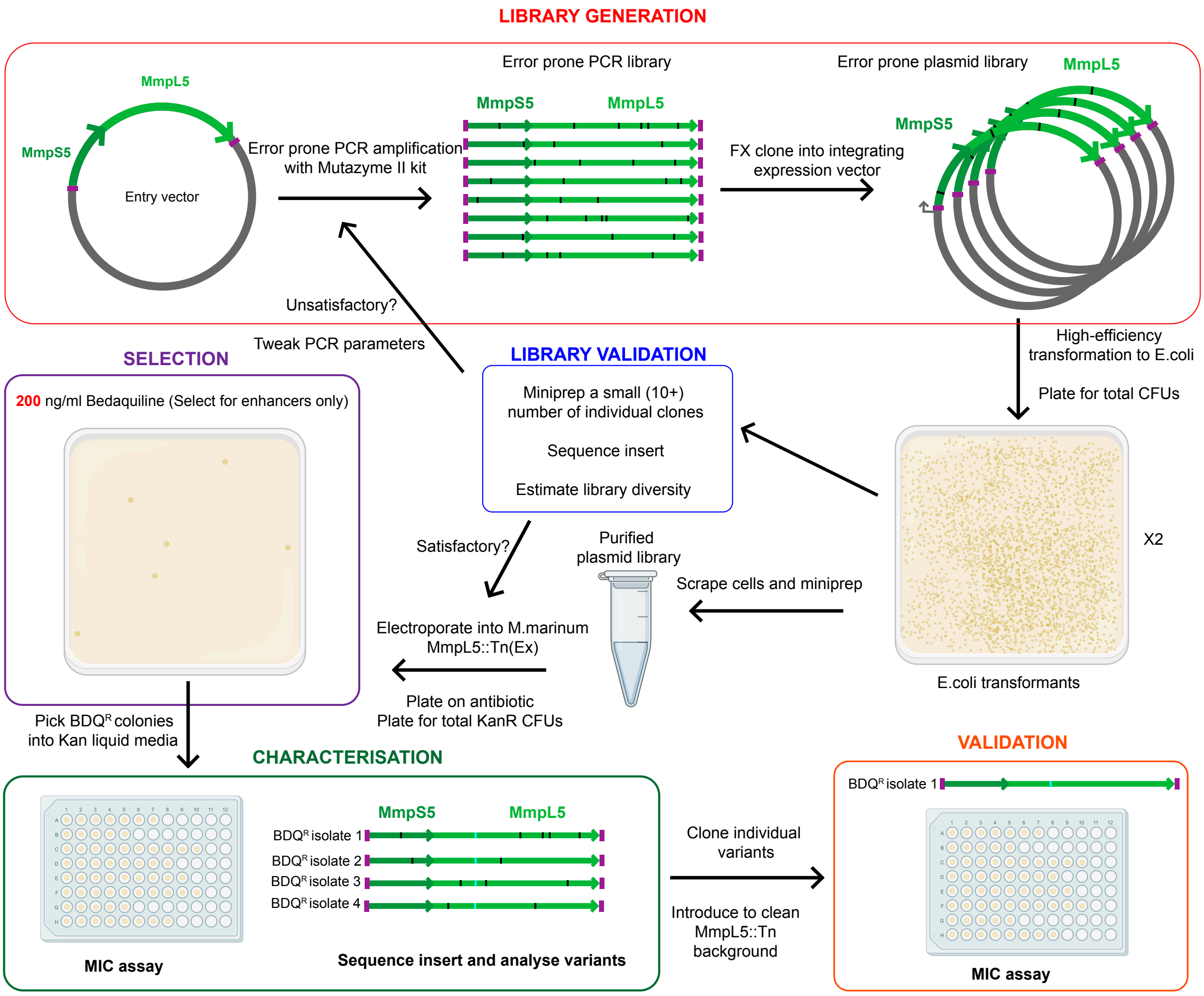

**Fig. S14. Error-prone PCR library generation scheme**

Outline of error-prone PCR library generation and selection experiments. In the library generation step, the coding sequence is PCR amplified from an FX-compatible entry vector and assembled into the integrative pMINTF3 vector. The library was electroporated into *E. coli*, and plasmid DNA from a subset of colonies was sequenced to estimate mutation rate. Library DNA was isolated from the plate.

#### Suppressor selection

| MmpL5 variant | CFU plated | Number resistant clones | Resistance Frequency |
|---|---|---|---|
| N444K | $2.78 \times 10^{9}$ | 12 | $2.3 \times 10^{-8}$ |

| Resistant isolate(s) | Genotype |
|---|---|
| Sup-1–Sup-4, Sup-8–Sup11 | Recombination MMAR_1005::Tn with $L5_{attB}$::pFLAG-attP-Kan *Mtb MmpS5L5* N444K |
| Sup-5 and 6 | $L5_{attB}$::pFLAG-attP-Kan *Mtb MmpS5L5* N444K, Y331D |
| Sup-7 | atpE A63P |
| Sup-12 | atpE A63V |

#### EP-PCR variant selection

| Bedaquiline concentration | Total Kan$^R$ colonies | BDQ$^R$,Kan$^R$ colonies | Proportion of library |
|---|---|---|---|
| 200 ng ml$^{-1}$ | $7.6 \times 10^{4}$ | 5 | 0.005% |

| Resistant isolate(s) | Number of mutations | Nucleotide/amino change |
|---|---|---|
| BDQ-2 | 6 | *MmpL5*: c.587T>A (p.Val193Asp), c.704T>A (p.Met235Lys), c.1269C>T (p.Ala423Ala), c.1279G>T (p.Ala427Ser), c.1447G>T (p.Ala483Ser), c.2527C>T (p.Leu843Leu) |
| BDQ-3 and BDQ-5 | 4 | *MmpS5*: c.122A>G (p.Lys42Arg) *MmpL5*: c.587T>A (p.Val193Asp), c.903C>G (Thr301Thr), c.2106A>G (Gln702Gln), |
| BDQ-6 | 5 | *MmpL5*: c.587T>A (p.Val193Asp), c.945G>A (p.Leu315Leu), c.1087C>A (p.Arg363Arg), c.1162C>T (Arg388Cys), c.1890C>T (p.Leu630Leu), |
| BDQ-7 | 4 | *MmpL5*: c. 755T>A (p.Phe252Tyr), c.1766A>G (p.Lys587Arg), c.2356C>A (p.Leu786Met), c.2705T>C (p.Val902Ala) |

**Fig. S15. Summary of resistance selection experiments**

Tables summarising the input parameters and outcomes of the suppressor selection (Upper) and EP-PCR variant selection experiments. Sup-# refers to the isolate from the suppressor selection. BDQ-# refers to the isolate from the EP-PCR variant selection.

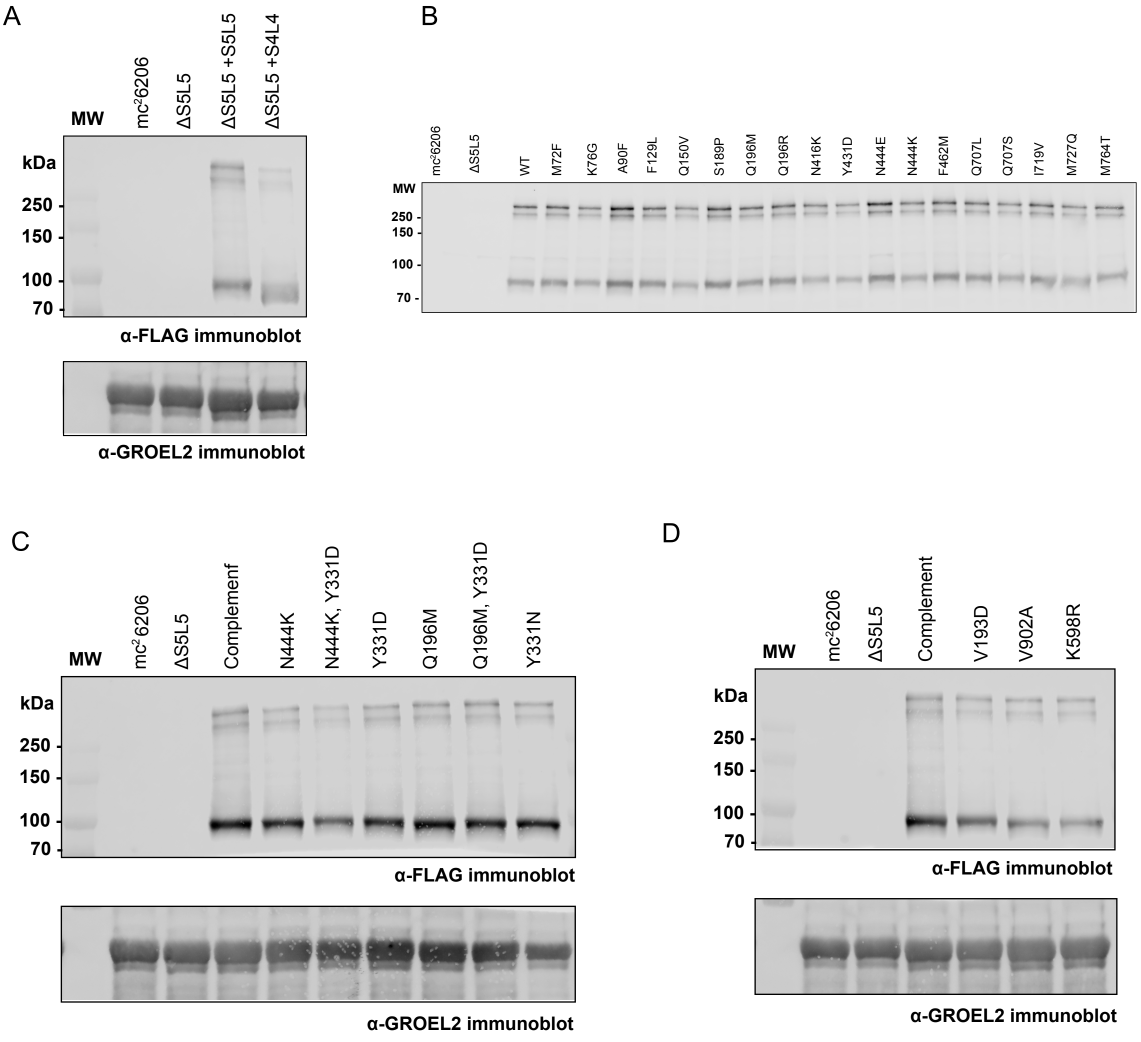

**Fig. S16. Immunoblot analysis of MmpS5L5 mutant strains**

(A–D) Immunoblot analysis of the strains used in this work. 20 μg total protein was loaded per lane. Upper blots indicate anti-FLAG immunoblot to detect FLAG-tagged MmpL5. Lower blots are loading controls against the cytoplasmic protein GROEL-2.

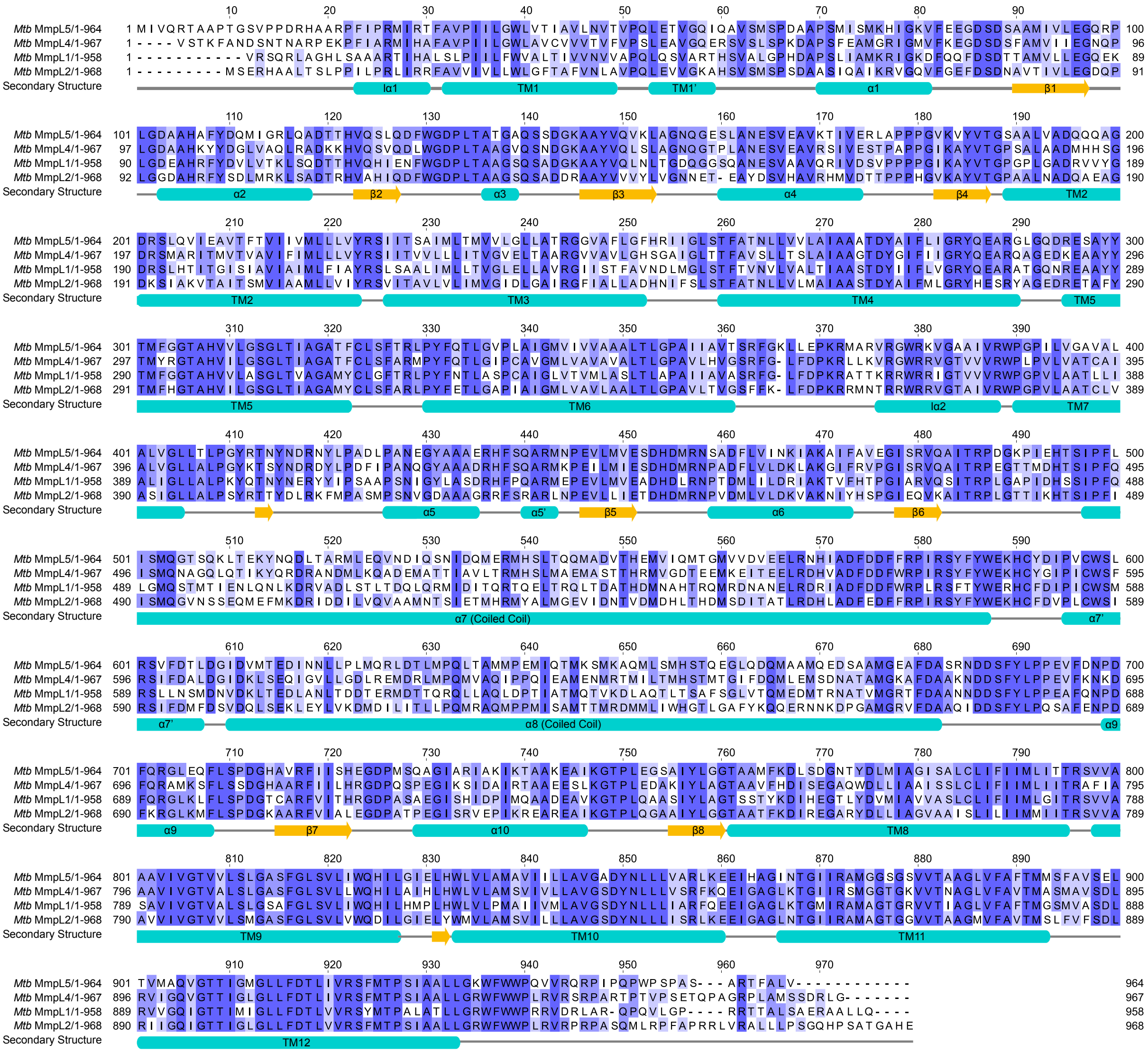

### Fig. S17. Multiple sequence alignment of MmpL5/4/2/1

Multiple sequence alignment of MmpL5/4/2/1. Numbering is according to position in MmpL5. Residues are coloured according to strength of sequence conservation. Secondary structure elements are indicated below the alignment.

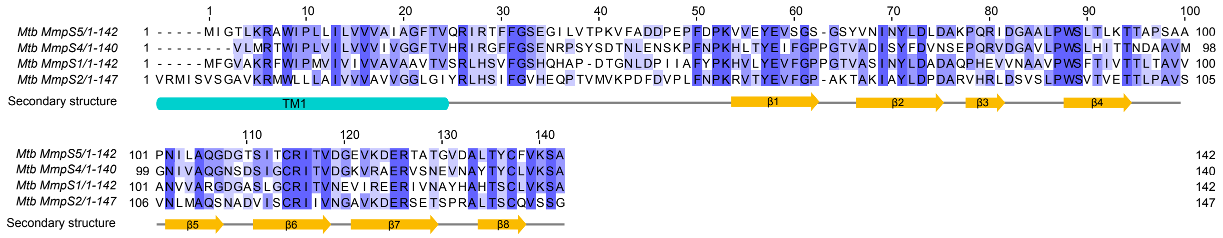

### Fig. S18. Multiple sequence alignment of MmpS5/4/2/1

Multiple sequence alignment of MmpS5/4/2/1. Numbering is according to position in MmpS5. Residues are coloured according to strength of sequence conservation. Secondary structure elements are indicated below the alignment.

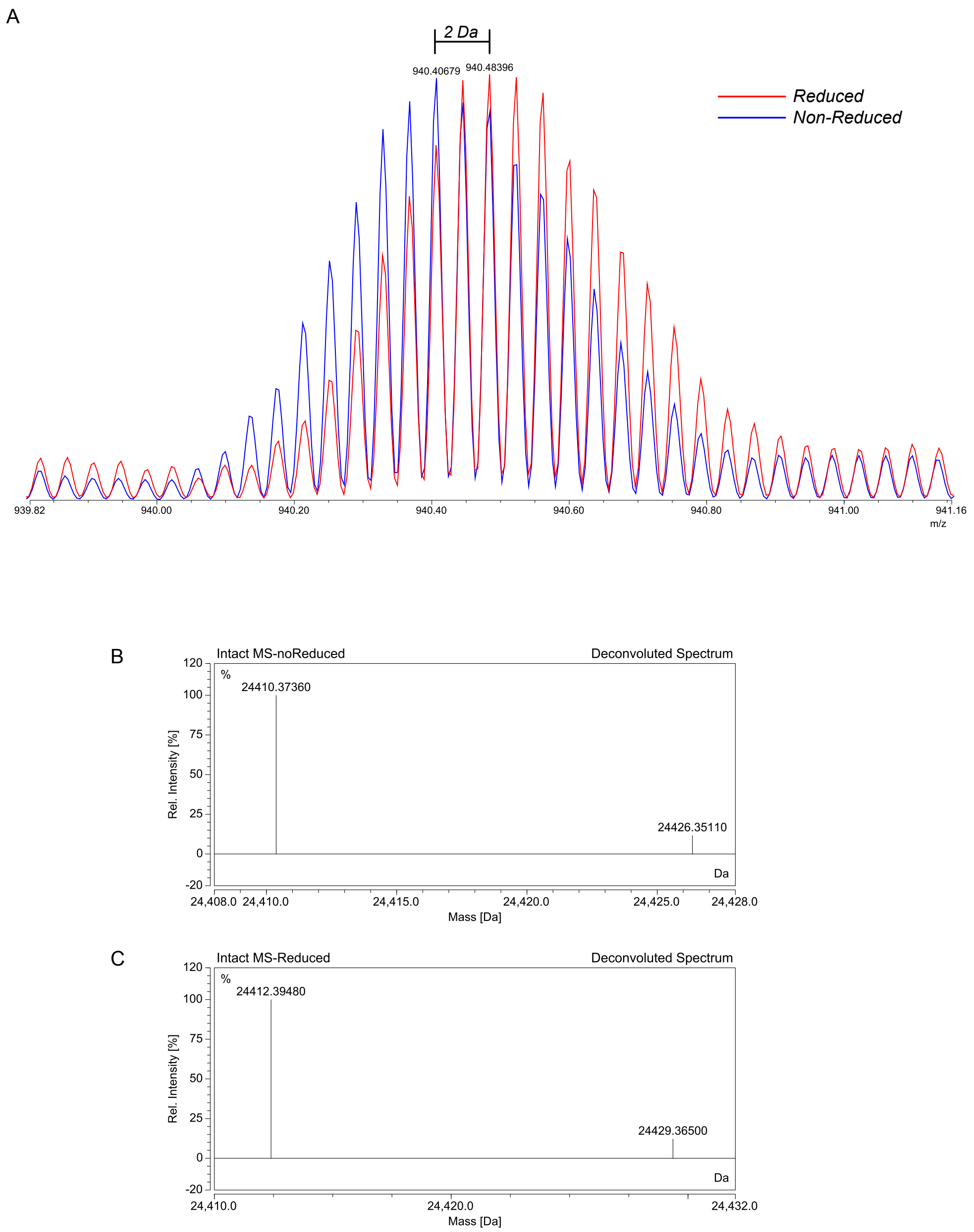

**Fig. S19. Mass spectra of purified MmpL5 coiled-coil domain (493–683) under non-reducing and reducing conditions.**

(A) Raw mass spectrometry spectra of the MmpL5 coiled-coil domain. The blue trace represents the non-reduced sample, showing a distinct isotopic envelope. The red trace corresponds to the TCEP-reduced sample (B) Deconvoluted mass spectrum of non-reduced MmpL5 coiled-coil domain**.** The primary peak at 24410.37 Da corresponds to the expected mass of the oxidized coiled-coil domain. (C) Deconvoluted mass spectrum of TCEP-reduced MmpL5 coiled-coil domain. The primary peak at 24412.39 Da, approximately 2 Da higher than the non-reduced sample, confirms the presence of an intramolecular disulfide bond that is reduced by TCEP.

### Table S1. Summary of bacterial strains and constructs used in this study

#### Bacterial strains

| Reagent or resource | Description | Source | Identifier |
|---|---|---|---|
| Organisms/Strains | | | |
| *E. coli* DH5α | | Invitrogen | |
| *M. smegmatis* mc$^2$155 | | Snapper *et al*., 1990 | |
| *M. tuberculosis* H37Rv mc$^2$6206 Δ*panCD* Δ*leuCD* | | Vilcheze et al., 2018 | |
| *M. tuberculosis* H37Rv mc$^2$6206 Δ*panCD* Δ*leuCD* Δ*S5L5*::loxP | Strain containing a deletion of the MmpS5L5 operon, generated by recombineering | Fountain and Waller *et al.* 2025 | |
| *M. tuberculosis* H37Rv mc$^2$6206 Δ*panCD* Δ*leuCD* Δ*S5L5*::loxP attB$_{L5}$::pMINTF3 *Mtb MmpS5L5* | Complemented ΔS5L5 strain with integrative pMINTF3 vector expressing *Mtb* MmpS5L5. | Fountain and Waller *et al.* 2025 | |
| *M. marinum* 'M' strain | | | |
| *M. marinum* MMAR::1005::Tn | | Ramakrishnan lab, unpublished | |
| *M. marinum* MMAR::1005::Tn attB$_{L5}$::pFLAG-attP-Kan *Mtb MmpS5L5* | | This study | |

#### DNA constructs

| Reagent or resource | Description | Source | Identifier |
|---|---|---|---|
| pMEXC3GF | ATc-inducible episomal expression construct with C-terminal GFP-FLAG tag | This study | |
| pMEXC3GF *Mtb MmpS5L5(*ΔCC) | ATc-inducible episomal expression construct encoding MmpS5 and MmpL5 ΔCC (Δ494–687) | This study | |
| pET-His6-TEV-MmpL5(492–683) | *E. coli* expression construct of *E. coli* codon-optimised MmpL5 coiled-coil domain | This study | |
| pINIT-Cat | FX cloning entry vector | Geertsma and Dutzler, 2011 | Addgene #46858 |
| pMA-Int | *E. coli* replicative vector which contains the mycophage L5 integrase gene required for attP/attB integration. | Arnold et al. 2018 | Addgene #110096 |
| pINIT-Cat *Mtb MmpS5L5* | FX cloning entry vector containing SapI-domesticated *Mtb* MmpS5L5 operon | This study | |
| pINIT-Cat *Mtb MmpS4L4* | FX cloning entry vector containing SapI-domesticated *Mtb* MmpS4L4 operon | This study | |
| pFLAG-attP-Kan | Single copy integrative vector, with L5 attP, but lacking L5 int. FX cloning compatible, with ccdB. Constitutive pmyctetO promoter and C-terminal 3x FLAG tag. | This study | |
| pFLAG-attP-Kan *Mtb-MmpS5L5* | pFLAG-attP-Kan containing a SapI-domesticated *Mtb* MmpS5L5 operon. | This study | |
| pFLAG-attP-Kan *Mtb MmpS5L5* N444K | | This study | |
| pMINTF3 | Single copy integrative vector, with mycobacteriophage L5 int and attP. FX cloning compatible, with ccdB. Constitutive pmyctetO promoter and C-terminal 3x FLAG tag. | This study | |
| pMINTF3 *Mtb MmpS5L5* | pMINTF3 containing a SapI-domesticated *Mtb* MmpS5L5 operon. | This study | |
| pMINTF3 *Mtb MmpL5* | pMINTF3 containing a SapI-domesticated *Mtb* MmpL5 | This study | |

| Reagent or resource | Description | Source | Identifier |
|---|---|---|---|
| DNA constructs | | | |
| pMINTF3 *Mtb MmpS1-MmpL5* | | This study | |
| pMINTF3 *Mtb MmpS2-MmpL5* | | This study | |
| pMINTF3 *Mtb MmpS4-MmpL5* | | This study | |
| pMINTF3 *Mtb MmpS5L5* Y277F | | This study | |
| pMINTF3 *Mtb MmpS5L5* D849N | | This study | |
| pMINTF3 *Mtb MmpS5L5* Y850F | | This study | |
| pMINTF3 *Mtb MmpS5L5 ∆CC* (494–687) | | This study | |
| pMINTF3 *Mtb MmpS5* K54A *MmpL5* | | This study | |
| pMINTF3 *Mtb MmpS5* K140A *MmpL5* | | This study | |
| pMINTF3 *Mtb MmpS5* K54A, K140A MmpL5 | | This study | |
| pMINTF3 *Mtb MmpS5* C137S *MmpL5* | | This study | |
| pMINTF3 *Mtb MmpS5L5* M72F | | This study | |
| pMINTF3 *Mtb MmpS5L5* K76G | | This study | |
| pMINTF3 *Mtb MmpS5L5* A90F | | This study | |
| pMINTF3 *Mtb MmpS5L5* F129L | | This study | |
| pMINTF3 *Mtb MmpS5L5* Q150V | | This study | |
| pMINTF3 *Mtb MmpS5L5* S189P | | This study | |
| pMINTF3 *Mtb MmpS5L5* Q196M | | This study | |
| pMINTF3 *Mtb MmpS5L5* Q196R | | This study | |
| pMINTF3 *Mtb MmpS5L5* D417L | | This study | |
| pMINTF3 *Mtb MmpS5L5* Y431D | | This study | |
| pMINTF3 *Mtb MmpS5L5* N444K | | This study | |
| pMINTF3 *Mtb MmpS5L5* N444E | | This study | |
| pMINTF3 *Mtb MmpS5L5* F462M | | This study | |
| pMINTF3 *Mtb MmpS5L5* Q707S | | This study | |
| pMINTF3 *Mtb MmpS5L5* Q707L | | This study | |
| pMINTF3 *Mtb MmpS5L5* I719V | | This study | |
| pMINTF3 *Mtb MmpS5L5* M727Q | | This study | |
| pMINTF3 *Mtb MmpS5L5* M764T | | This study | |
| pMINTF3 *Mtb MmpS5L5* H832Y | | This study | |
| pMINTF3 *Mtb MmpS5L5* N444K, Y331D | | This study | |
| pMINTF3 *Mtb MmpS5L5* Y331D | | This study | |
| pMINTF3 *Mtb MmpS5L5* Y331N | | This study | |
| pMINTF3 *Mtb MmpS5L5* V193D | | This study | |
| pMINTF3 *Mtb MmpS5L5* V902A | | This study | |
| pMINTF3 *Mtb MmpS5L5* K598R | | This study | |
| pMINTF3 *Mtb* MmpS4L4 | Construct expressing MmpS4L4 | This study | |

### Table S2. Oligonucleotide sequences used in this study

| Oligo | Sequence | Description |
|---|---|---|
| P104 | ATATATGCTCTTCTAGTCTAATGCGGACTTGGATTCCACTGGTCATCCTGG | Amplification of MmpS4L4 for insertion into FX-compatible vectors |
| P105 | TATATAGCTCTTCATGCGCCGAGGCGGTCGCTGCTCATC | |
| P106 | ATATATGCTCTTCTAGTATTGGAACTCTCAAGCGTGCCTG | Amplification of MmpS5L5 for insertion into FX-compatible vectors |
| P107 | TATATAGCTCTTCATGCGACCAAGGCGAAGGTCCGTG | |
| P298 | CGGACCTCTATTCACAGGGTACG | For constructing pFLAG-attP-Kan |
| P299 | GCTGGAAGCACTCAACCTCG | |
| P300 | GTACCCTGTGAATAGAGGTCCGGTGTCTCAAAATCTCTGATGTTAC | |
| P301 | CGAGGTTGAGTGCTTCCAGCTCCTTCAACTCAGCAAAAG | |
| P352 | TCCATGAAGGCGCAGATGCTG | For SapI domestication of the MmpS5L5 sequence |
| P353 | CTTCATGGTCTGGATCATCTCGG | |
| P354 | GCCTGCTGTTCGACACCCTGATC | |
| P355 | CCATACCGATGGTGGTGCCAAC | |
| P366 | ATCGTGCAAAGGACAGCTGC | For deletion of MmpS5 to create MmpL5 only construct |
| P367 | ACTAGAAGAGCGGCCAACCAG | |
| P368 | CTCGATCGGTTTGCCGTCCG | Use to remove MmpL5' coiled coil domain |
| P369 | TCGTTCTATCTGCCTCCCGAGGTTTTC | |
| P465 | TTACTTGTCATCGTCGTCCTTGTAATCACTAGTTTTGTAGAGCTCATCCATGCC | For constructing pMEXC3GF vector |
| P466 | CATCATTGAGTTTAAACGGTCTCCAGC | |
| P577 | CGGCCTGGTTGGCCGCTCTTCTAGTTTCGGCGTTGCCAAACGCTTC | PCR of MmpS1 with overhangs for replacing MmpS5 |
| P578 | TCGGCGCAGCTGTCCTTTGCACGATCATGCGGATTTCACCAGGCAACTG | |
| P579 | CGGCCTGGTTGGCCGCTCTTCTAGTAGGATGATTTCGGTTAGCGGC | PCR of MmpS2 overhangs for replacing MmpS5 |
| P580 | CGTCGGCGCAGCTGTCCTTTGCACGATCATCCGGATGACACCTGGC | |
| P581 | CGGCCTGGTTGGCCGCTCTTCTAGTCTAATGCGGACTTGGATTCCACTG | PCR of MmpS4 with overhangs for replacing MmpS5 |
| P582 | TCGGCGCAGCTGTCCTTTGCACGATCATGCGGACTTCACCAAGC | |
| P583 | ATGATCGTGCAAAGGACAGCTG | For replacing MmpS5 with paralog. Used with P367 |
| P623 | GCGTTTTACGACCAGATGATCGG | Amplify MmpS5L5 without residues 2–106 for assembly with eblock |
| P624 | TGCGGATTTCACAAAGCAGTAGGTC | |
| P625 | GATCGAGGCGGTCACGTTCAC | Amplify MmpS5L5 without residues 115–205 for assembly with eblock |
| P626 | CGATCATCTGGTCGTAAAACGCATGG | |
| P627 | GTACTACACCATGTTCGGCGG | Amplify MmpS5L5 without residues 207–297 for assembly with eblock |
| P628 | CACCTGCAGACTACGGTCG | |
| P629 | CTACCGGACCAACTACAACGACCG | Amplify MmpS5L5 without residues 298–409 for assembly with eblock |
| P630 | CGACTCCCGGTCCTGGCC | |
| P631 | GAAACTGACCGAGAAATACAACCAGGAC | Amplify MmpS5L5 without residues 410–507 for assembly with eblock |
| P632 | GGGTCAGCAGACCGACGAG | |
| P633 | GTCATGACCGAAGACATCAACAACCTG | Amplify MmpS5L5 without residues 508–611 for assembly with eblock |
| P634 | CTGGTGCCCTGCATGCTGATC | |
| P635 | GGCCTGGAACAGTTCCTCTCG | Amplify MmpS5L5 without residues 612–703 for assembly with eblock |
| P636 | GTCGATTCCGTCGAGGGTGTC | |
| P637 | TTCATCATCATGCTGATCACCACCC | Amplify MmpS5L5 without residues 704–787 for assembly with eblock |
| P638 | GCGTTGGAAGTCGGGATTGTC | |
| P639 | GGCATCATCCGTGCGATGG | Amplify MmpS5L5 without residues 788–868 for assembly with eblock |
| P640 | GATCAGGCAGAGTGCGGAGATTC | |
| P641 | GCATGAAGAGCGGCCACC | Amplify MmpS5L5 without residues 869–964 for assembly with eblock |
| P642 | GGTGTTGATTCCGGCGTGGATC | |
| P677 | GTTCTCGGCTCGATGATCCACC | For sequencing MmpS5L5 from integrated constructs |
| P678 | GCGAGTCAGTGAGCGAGGAAG | |
| P721 | GCAACTCTTGTGCGACTCTTCTGAC | For construction of pMINTF3 |
| P722 | CGGTTCCTGGCCTTTTGCTG | |
| P723 | ACATGTGAGCAAAAGGCCAGC | |
| P724 | CGTATGCCCAGGTCAGAAGAGTC | |
| P725 | GCCTGGTTGGCCGCTCTTCTAGTATTGG | For mutagenic PCR of MmpS5L5 with Mutazyme II |
| P726 | CCTCGGTGGCCGCTCTTCATGC | |
| P812 | ATTGTAAAACGACGGCCAGTAAAGAAAAACGTCCCGTTGGGAAC | PCR of *Mm atpE* for sequencing with M13F |
| P813 | TTAAAATCCCGACCCCATACCAG | |

### Table S3. Cryo-EM data collection, refinement, and validation statistics

| **Data collection and processing** | **MmpS5L5-BDQ** [EMDB-53947, PDB 9RGB] | **MmpS5L5 (apo)** [EMDB- 53941, PDB 9RFU] |
|---|---|---|
| Microscope | Krios Titan G4 | Krios Titan G4 |
| Magnification | 130,000 | 130,000 |
| Voltage (kV) | 300 | 300 |
| Electron exposure ($e^{-}$/Å$^2$) | 80 | 80 |
| Defocus range (μm) | -0.6 to -2.2 | -0.6 to -2.2 |
| Pixel size (Å) | 0.955 | 0.955 |
| Symmetry imposed | C1 | C1 |
| Initial particle images (no.) | 3,240,490 | 4,810,392 |
| Final particle images (no.) | 33,185 | 27,426 |
| Map resolution (Å) | 3.1 | 3.3 |
| FSC threshold | 0.143 | 0.143 |
| Map resolution range (Å) | 2.7–3.7 | 2.9–3.8 |
| **Model Refinement** | | |
| Initial model used | AlphaFold2 | AlphaFold2 |
| Map sharpening *B* factor (Å$^2$) | -54.7 | -65.2 |
| Model composition | | |
| Non-hydrogen atoms | 19410 | 19410 |
| Protein residues | 2543 | 2543 |
| Ligands | 6 | 6 |
| R.m.s. deviations | | |
| Bond lengths (Å) | 0.005 | 0.005 |
| Bond angles (°) | 0.908 | 1.067 |
| Validation | | |
| MolProbity score | 1.15 | 1.62 |
| Clashscore | 3.64 | 7.85 |
| Poor rotamers (%) | 0.2 | 0 |
| Ramachandran plot | | |
| Favored (%) | 98.10 | 96.83 |
| Allowed (%) | 1.86 | 3.17 |
| Disallowed (%) | 0.04 | 0 |
